# Functional Characterization of the Lin28/let-7 Circuit during Forelimb Regeneration in *Ambystoma mexicanum* and its Influence on Metabolic Reprogramming

**DOI:** 10.1101/2020.06.18.160291

**Authors:** Hugo Varela-Rodríguez, Diana G. Abella-Quintana, Luis Varela-Rodríguez, David Gomez-Zepeda, Annie Espinal-Centeno, Juan Caballero-Pérez, José Juan Ordaz-Ortiz, Alfredo Cruz-Ramírez

## Abstract

The axolotl (*Ambystoma mexicanum*) is a caudate amphibian, which has an extraordinary ability to restore a wide variety of damaged structures by a process denominated epimorphosis. While the origin and potentiality of progenitor cells that take part during epimorphic regeneration are known to some extent, the metabolic changes experienced and their associated implications, remain unexplored. However, a circuit with a potential role as a modulator of cellular metabolism along regeneration is that formed by Lin28/let-7. In this study, we report two Lin28 paralogs and eight mature let-7 microRNAs encoded in the axolotl genome. Particularly, in the proliferative blastema stage amxLin28B is more abundant in the nuclei of blastemal cells, while the microRNAs amx-let-7c and amx-let-7a are most downregulated. Functional inhibition of Lin28 factors increase the levels of most mature let-7 microRNAs, consistent with an increment of intermediary metabolites of the Krebs cycle, and phenotypic alterations in the outgrowth of the blastema. In summary, we describe the primary components of the Lin28/let-7 circuit and their function during axolotl regeneration, acting upstream of metabolic reprogramming events.

## INTRODUCTION

Regeneration is a biological phenomenon that allows the restoration of tissues and biological structures, which have been damaged or lost in the organisms (Gilbert, 2010). While most eukaryotic organisms display certain ability to regenerate, such capacity varies along the animal phylogeny, and relies on specific strategies according to the origin and potentiality of the cells that drive the process (Bely and Nyberg, 2010). Epimorphic regeneration in salamanders is a major regenerative strategy that allows the recovery of complex biological structures, normally recalcitrant to regenerate in other tetrapods, such as mammals (Roy and Gatien, 2008). The main distinctive of epimorphosis is the establishment of a heterogeneous population of cells called blastema, originated in principle by reactivation of tissue-resident stem cells and dedifferentiation of mature cells, which will give rise to a new functional structure through later morphogenetic events (Leigh et al., 2018).

Since epimorphosis depends on dynamical changes in the cellular environment, with rapid transitions between states of proliferation and differentiation, it results interesting to unravel the functional roles of molecular circuits with potential to finely modulate such cell reprogramming events (Stocum, 2017). A promising circuit is the one conformed by Lin28 proteins and let-7 microRNAs (miRNAs), highly conserved among bilaterians (Roush and Slack, 2008). It has been reported that such circuit plays a key role as a regulator of the terminal cell differentiation in mammals, maintaining the cell fate, through coordination of cell proliferation, growth, and energy utilization (Thornton and Gregory, 2012). The function of Lin28 has been described in diverse animal models, as RNA-binding proteins that interact with precursors of the let-7 family, negatively regulating the abundance of mature let-7 microRNAs (Nowak et al., 2017). Alternatively, it has been demonstrated that Lin28 factors modulate the translation of diverse mRNAs (Mayr and Heinemann, 2013). In mammals, some Lin28 target transcripts are implicated in the cell cycle entry (*Ccn-a/b/d* and *Cdk-1/2/4/6*), chromatin remodeling (*Hmg-a1*), alternative splicing of mRNAs (*hnRnp-f, Tia-1* and *Tdp-43*), or primary metabolism (*Hk-1, Pfk-p* and *Pdk-1*) (Balzeau et al., 2017; Wilbert et al., 2012). Although the functional role of the Lin28 family members has been widely described as proliferative enhancers in oncogenesis (Nguyen et al., 2014), reprogramming factors of cell metabolism (Zhang et al., 2016; Zhu et al., 2011), and molecular timers during developmental transitions in organogenesis and metamorphosis (Faunes et al., 2017; Yermalovich et al., 2019), these proteins have not been sufficiently studied in the context of regeneration.

Pioneer studies have related some components of the Lin28/let-7 circuit with regenerative processes, as those that report a high regenerative plasticity in juvenile stages of *Caenorhabditis elegans*, where immature neurons with low levels of mature let-7 retain a robust regeneration at the axon disruption site, nearby to neural body (Nix and Bastiani, 2013; Zou et al., 2013). Similarly, the transient overexpression of Lin28 in postnatal sensory neurons of mouse after injury, induce an axonal regeneration *in vivo* through changes in the balance of the AKT-mTOR pathway (Wang et al., 2018). Therefore, an adequate cell metabolic state is relevant for regeneration, since the AKT-mTOR pathway acts as an important mediator between anabolic and catabolic cell reactions (Altomare and Khaled, 2012; Saxton and Sabatini, 2017). In this sense, it has been shown that the inducible overexpression of Lin28A in mouse neonatal tissues, improves regeneration by a rewiring of the primary energetic metabolism, where the glycolysis is favored to increase intermediary metabolites of the Krebs cycle (Shyh-Chang et al., 2013). However, such metabolic reprogramming of the cell bioenergetics differs from other metabolic profiles also achieved with overexpression of Lin28A, but in the context of embryonic development, or during active proliferation of primed pluripotent stem cells and malignant neoplastic cells (Ma et al., 2014; Miyazawa et al., 2017; Zhang et al., 2016).

Since the role of the Lin28/let-7 circuit has not been directly studied in the context of epimorphosis, using an amphibian model with a high and innate regenerative capacity, we decided to characterize its behavior and function during forelimb regeneration in axolotl (*Ambystoma mexicanum*). In this study, we describe the spatio-temporal expression dynamics of Lin28 proteins and changes in abundance of the let-7 microRNAs along diverse regeneration stages. Also, we implemented different metabolomics approaches to profile the metabolic variations that occur during epimorphic regeneration. Finally, we demonstrate that functional inhibition of Lin28 factors in key stages of limb regeneration, alters the regenerative process with a rewiring of the primary cell metabolism.

## MATERIALS AND METHODS

### Care and Handling of Animals

The axolotls were housed in a bioterium designed for the maintenance of their colonies, being obtained from a Unit of Environmental Management for *ex situ* reproduction at the Centro de Investigaciones Acuáticas de Cuemanco (UAM Campus Xochimilco). The animals were preserved individually at 17 °C in tap water with 12:12 hours light/dark cycle, using healthy late juvenile axolotls with a snout-tail length of 14-16 cm and a body mass range of 25-29 g, carefully amputated for first time in their right forelimbs at middle-zeugopod level. All surgical procedures were performed in specimens previously anesthetized with 0.02% benzocaine in swimming water.

### Pharmacological Treatments Application

Functional inhibition of Lin28 proteins was assayed employing the compound Lin28-1632 (*N*-Methyl-*N*-[3-(3-methyl[1,2,4]triazolo[4,3-*b*]pyridazin-6-yl)phenyl]acetamide; CAS number 108825-65-6) (Tocris Bioscience). The inhibitor was topically delivered by immersion of limbs into a solution at 80 µM of Lin28-1632, 0.75X PBS, 0.002% DMSO. In a first assay, the inhibitor was applied following a periodic scheme by 1 hour of topical administration daily for 6 consecutive days before the sample collections, at 6, 14, 20, and 42 days post amputation (dpa) (Fig. 5A). In a second assay, the inhibitor was applied continuously by 1 hour of topical administration every day, starting at 24 hours after amputation until 24 dpa, collecting the samples at 56 dpa (Fig. S2A). The pharmacological assays effectuated *in vivo* were accomplished with anesthetized axolotls kept on a wet bed with swimming water. The assays were carried out with three biological replicates for Lin28-1632 applications and two controls implemented, one control with topical administration of vehicle only and another without any treatment.

### Bioinformatics Analyses

The axolotl *lin-28* transcripts were obtained extending a previously reported transcriptome (Caballero-Pérez et al., 2018) with a pooled mRNA-seq library during axolotl embryo development at 4, 8, 9, 10, 12, 18, 19, 25, 30, 39 stages (unpublished data). New transcriptome was assembled with Trinity v2.0.6 (Grabherr et al., 2011), and contigs merged with CD-HIT-EST v4.6 (Fu et al., 2012) allowing at least 95% identity with 95% coverage. The axolotl *lin-28* transcripts were searched using human *LIN28A* and *LIN28B* sequences as references through Blast+ v2.2.30 (Camacho et al., 2009), keeping best matches with E-value <1×10e-5. The translated Lin28 cistrons were aligned using Geneious v8.1.9 (Kearse et al., 2012). Afterwards, domain architecture identification for axolotl Lin28 proteins was done using normal SMART v8.0 (Letunic and Bork, 2017), and the results graphed with IBS v1.0.3 (Liu et al., 2015). The phylogenetic tree for the Lin28 family in vertebrates was made with *lin-28* coding sequences aligned employing DECIPHER v1.12.1 (Wright, 2015). Next, the best-fit model of nucleotide substitution was determined with PartitionFinder v2.1.1 (Lanfear et al., 2016), setting a linked branch length and the “greedy” heuristic search algorithm with corrected Akaike Information Criterion, defining each codon position for each domain of Lin28 as data blocks. The partitioned phylogenetic analysis was realized with IQ-Tree v1.3.11.1 (Nguyen et al. 2014), establishing the *lin-28* sequence of *Branchiostoma belcheri* as outgroup, under the parameters “-bb 10000 -bspec GENESITE -allnni -seed 12345”. The phylogenetic tree inferred was annotated and graphed with iTOL v4.4.2 (Letunic and Bork, 2019), indicating the accession numbers of the *lin-28* sequences obtained from NCBI. Some *lin-28* transcripts were acquired from public transcriptomes, being the case for *Pleurodeles waltl* (Matsunami et al., 2019) and *Cynops pyrrhogaster* (Japan Newt Research Community, 2013). New *lin-28* transcripts generated were submitted at NCBI and will be available upon manuscript acceptance.

Identification of axolotl let-7 microRNAs was performed with a raw sRNA-seq dataset of *A. mexicanum* from the SRA study SRP093628 at NCBI (Caballero-Pérez et al., 2018). The quality of sequencing data was checked using FastQC v0.11.6 (Andrews, 2010), and sequencing adapters removed with cutadapt v1.7.1 (Martin, 2011) under the parameters “-e 0.1 -O 5 -m 15” running on GNU Parallel v20161222 (Tange, 2011). Subsequently, low-quality reads were filtered through Sickle v1.33 (Joshi and Fass, 2011) with the parameters “-q 30 -l 15”. Afterwards, sequencing reads were mapped with Geneious v8.1.9 (Kearse et al., 2012) through the criteria of 0% mismatches, 22 word length, without gaps, using the mature let-7 sequences reported in the miRBase database v21.0 (Griffiths-Jones et al., 2006) as reference.

The primary let-7 transcripts of axolotl were reconstructed with a raw mRNA-seq dataset of *A. mexicanum* from the SRA study SRP093628 at NCBI (Caballero-Pérez et al., 2018). The quality of sequencing data was evaluated with FastQC v0.11.6 (Andrews, 2010), and both adapters as low-quality reads were detached with Trimmomatic v0.36 (Bolger et al., 2014) under the parameters “SLIDINGWINDOW:4:20 MINLEN:35”. The transcripts reconstruction was realized with Geneious v8.1.9 (Kearse et al., 2012). The axolotl let-7 microRNAs were used as reference templates for transcript reconstructions. The transcripts were confirmed in the axolotl genome v3.0 (Nowoshilow et al., 2018), and the results graphed with IBS v1.0.3 (Liu et al., 2015). The primary let-7 sequences reconstructed were submitted will be available at NCBI after manuscript acceptance.

The let-7 precursors encoded in the primary transcripts of axolotl were searched with Rfam v14.1 (Kalvari et al., 2018). A consensus model was generated to represent the secondary structures predicted for let-7 precursors with RNAalifold v2.4.8 (Bernhart et al., 2008) under the parameters “--mis -p -r -d2 --noClosingGU -T 20”, using an alignment generated with Geneious v8.1.9 (Kearse et al., 2012). The provided model was plotted with R-chie (Lai et al., 2012).

### Transcriptional Expression Measurements by qRT-PCR

Total RNA from different regeneration stages was isolated of freshly amputated samples (1 mm of distal-most tissue) rapidly frozen into liquid nitrogen, pulverized and then homogenized mechanically with a homogenizer BDC-2002 (Caframo) using TRIzol reagent (Invitrogen), according to manufacturer instructions. The quantity and purity of RNA extracted were evaluated with a NanoDrop 2000 (NanoDrop Technologies), and the integrity with a native agarose gel electrophoresis (Aranda et al., 2012). The cDNA synthesis from mRNA was performed using SuperScript III Reverse Transcriptase (Invitrogen) with 2 µg of total RNA by sample and oligo(dT)_20_, while the generation of cDNA from sRNA was done utilizing the Universal cDNA synthesis kit II (Exiqon) with 0.4 µg of total RNA by sample, in both cases following manufacturer protocols. The cDNA templates produced from mRNA and sRNA were diluted to 1:3 and 1:12 respectively, and then amplified into a real-time PCR system CFX96 (Bio-rad) using 2 µL of diluted cDNA with standard SYBR GREEN PCR master mix (applied Biosystems), following manufacturer procedures. The primer sequences employed are listed in Table S1. The data analyses were performed using GenEx v6.1.0.757 (MultiD Analyses AB), and the relative abundance of transcripts were determined by efficiency corrected -ΔCq and 2^-ΔΔCq^ (Fold Change) methods (Livak and Schmittgen, 2001). The normalization of transcripts abundances was done with two stable endogenous references, *odc-1* and miR-200b-5p, previously reported in axolotl transcripts measurements during limb epimorphosis for mRNAs and microRNAs respectively (Guelke et al., 2015; Holman et al., 2012). The qRT-PCR assays were performed with three biological replicates, each of them being independently measured three times for technical reproducibility.

### Fluorescent Immunolocalization Assays

Limbs from diverse regeneration stages were collected (3 mm of distal-most tissue), and rapidly fixed by immersion in a cold freshly prepared solution of 4% paraformaldehyde 0.75X PBS for 16-18 hours at 4 °C. Next, samples were rinsed 3 times for 10 minutes each with 0.75X PBS, and transferred to a 20% sucrose 0.75X PBS solution for 24 hours at 4 °C. Afterwards, limbs were embedded into Tissue-Tek O.C.T compound (Sakura Finetek), frozen with liquid nitrogen and preserved at −80 °C. Tissue sectioning was performed using a cryostat microtome Leica CM1860 at 10 µm of thickness, and sections mounted on gelatin coated slides then stored at −80 °C. For indirect immunofluorescence staining, sections were air dried 1 hour at 37 °C and treated 2 times for 10 minutes each with 0.3M glycine 1X PBS at room temperature. Subsequently, samples were permeabilized with 0.25% Triton X-100 1X PBS for 10 minutes and washed 3 times for 5 minutes each using 1X PBS. Next, sections were blocked with a solution of 5% goat normal serum 1X PBS 0.05% Tween-20 for 30 minutes at room temperature into a humid chamber. The primary antibodies used were a rabbit anti-Lin28A (ab170402, Abcam) diluted 1/200, a rabbit anti-Lin28B (HPA061745, Sigma-Aldrich) diluted 1/200, and a mouse anti-H3S10ph (ab14955, Abcam) diluted 1/1000, using blocking solution for antibody dilutions. Then, samples were incubated with primary antibodies for 16-18 hours at 4 °C into a humid chamber and rinsed 3 times for 10 minutes each with 1X PBS. Later, sections were incubated for 2 hours into a humid chamber with goat secondary antibodies, anti-mouse IgG Alexa Fluor 594 (ab150116, Abcam) or anti-rabbit IgG Alexa Fluor 488 (A-11008, Thermo Fisher Scientific), all secondary antibodies diluted 1/1000 in blocking solution at 25 °C, being then rinsed 3 times for 10 minutes each with 1X PBS. Samples were counterstained using DAPI at 1 µg/mL (Novus Biologicals) or TRITC-conjugated Phalloidin at 2 µg/mL (Sigma-Aldrich) in 1X PBS for 20 minutes into a humid chamber at 25 °C, and rinsed 3 times for 10 minutes each with 1X PBS. Finally, sections were mounted with Fluoroshield aqueous mounting medium for histology (Sigma-Aldrich) and stored at 4 °C in darkness. The fluorescent immunolocalization assays were performed with three biological replicates and two technical replicates for each of them.

### Hematoxylin & Eosin Staining

These dyes stain the nuclei in blue-purple, and the cytoplasm with different shades of pink (Varela-Rodríguez et al., 2020). Tissue sections were first post-fixed with 10% paraformaldehyde for 5 minutes and rinsed with tap water. Next, samples were stained with Gill Hematoxylin #1 (Sigma-Aldrich) for 10 minutes and washed with tap water. Then, slides were treated with acid ethanol (5% HCl in 70% ethanol) for 5 seconds and rinsed twice with distilled water. Afterwards, samples were immersed into ammoniacal water (0.4% hydroxide ammonium) for 5 minutes and rinsed with distilled water. Later, sections were stained with alcoholic Eosin Y (Sigma-Aldrich) for 30 seconds and washed with absolute ethanol. Finally, samples were cleared using 1:1 ethanol–xylene for 2 minutes, and 3 subsequent rounds of xylene for 5 minutes each one. The sections were mounted with DPX mountant medium (Sigma). Histological staining was performed with three biological replicates and two technical replicates for each of them.

### Iron Hematoxylin–Safranin–Fast Green–Aniline Blue Staining

These dyes stain the glycosaminoglycans and proteoglycans with shades of reddish orange, collagenous fibrils in dark blue, cell nuclei of black hue, and erythrocytes or Leydig cells in green-bluish. Tissue sections were processed similarly to an H&E staining. Iron Hematoxylin was prepared just before use by addition of acid ferric chloride (2% ferric chloride, 35 mM HCl) to the Gill Hematoxylin #1 (Sigma-Aldrich) in 1:10 proportion, respectively. After the ammoniacal water step in modified H&E, samples were stained with 0.1% Safranin O (Sigma-Aldrich) for 5 minutes and washed with distilled water. Then, sections were stained with 0.01% Fast Green FCF (Sigma-Aldrich) for 10 minutes and washed with distilled water. Afterwards, slides were immersed in phosphotungstic acid (1% phosphotungstic acid) for 1 minutes and rinsed with distilled water. Next, samples were stained with 0.5% Aniline Blue (Jalmek) for 1 minute and washed twice with distilled water. A differentiation step was performed, washing the slides rapidly with 1% acetic acid and rinsed with distilled water. Finally, samples were immersed in absolute ethanol for 5 seconds and cleared using 1:1 ethanol–xylene for 2 minutes, followed by 3 consecutive rounds with xylene for 5 minutes each one. The sections were mounted with DPX mountant medium (Sigma). Histological staining was performed with three biological replicates and two technical replicates for each of them.

### Microscopy and Image Processing

Macroscopic morphology was inspected with a Leica EZ4 stereoscopic microscope (Leica Microsystems). Histological phenotype was examined using a Leica DM6000B microscope (Leica Microsystems) with transmitted light and differential interference contrast, as well as a digital microscope VHX-5000 (Keyence) with reflected light in brightfield. A confocal Zeiss LSM800 with Axio Imager.Z2 (Carl Zeiss Microscopy) was employed for immunolocalizations, utilizing laser lines at 405, 488 and 561 nm. Quantitative measurements were performed using Fiji/ImageJ v2.0/1.52i (Schindelin et al., 2012) to determine signal colocalizations (Arqués et al., 2012) and surface areas. Statistical analyses were made with Minitab v16.1 (Minitab Inc.).

### Metabolite Extractions and Metabolomics Analyses

Limb samples from different regeneration stages were collected (1 mm of distal-most tissue), quickly washed 3 times with 0.75X PBS, and then grounded with liquid nitrogen. Next, pulverized samples were homogenized in cold methanol-water at 4:1 ratio respectively with 0.1% formic acid, and sonicated by 5 cycles of 10 seconds ON / 50 seconds OFF at 40% amplitude on ice bath utilizing a Branson Sonifier 150 sonicator (Emerson Electric) with microprobe of 1/8” thickness. Afterwards, metabolic extracts were incubated for 2 hours with shaking to 1,400 rpm at 4 °C, and then centrifuged 3 times for 10 minutes each to 14,000 rpm at 4 °C, collecting supernatants every time for further precipitate proteins and debris. Subsequently, supernatants were vacuum dried at 30 °C in a Genevac miVac centrifugal concentrator (SP Scientific) and kept at −80 °C.

Non-targeted and targeted metabolomics analyses were performed using ultra-high liquid chromatography coupled to high definition mass spectrometry (UPLC-HDMS). Chromatographic separation was achieved on an Acquity UPLC class I system (Waters Corporation), with an HSS-T3 C18 analytical column (2.1 × 100 mm, 1.8 µm particle size, Waters Corporation) maintained at 40 °C, using the respective chromatographic methods described below. The mobile phases consisted of A: deionized water containing 0.1% formic acid; and B: acetonitrile containing 0.1% formic acid. Data was acquired in an orthogonal QTOF Synapt HDMS G1 (Waters Corporation) with infusion of Leucine-enkephalin (2 ng/mL) at 5 μL/min for mass calibration (Lockmass). The UPLC-HDMS system was controlled using MassLynx v4.1 (Waters Corporation). Samples were stored at 4°C during analysis. All solvents used were HPLC grade (TEDIA), and water was Milli-Q grade (Merck KGaA). Raw data will be available at MetaboLights (Haug et al., 2020) after paper acceptance. Analytical standards (purity ≥99%) for citric, α-ketoglutaric, succinic, and fumaric acids, as well as formic acid (purity ≥98%), were purchased from Sigma-Aldrich.

For non-targeted metabolite profiling, dried extracts were reconstituted in a mixture of 1:1 methanol-water with 0.1% formic acid, using 5 µL of solvent per milligram of fresh tissue and 10 µL of each sample were injected. The non-targeted profiling and targeted quantification were performed with three biological replicates, and three quality controls generated by mixing a fixed portion of each sample analyzed. The compounds were eluted at 0.5 mL/min, starting at 5% B for 1 minute, followed by a linear gradient increase from 80% to 100% B in 10 min, a wash at 100% B during 1 min, and a re-equilibration at 5% B for 2 min. The HDMS was operated under in W mode (resolution of 17,000 FWHM). Data were continuously acquired using MS^E^ acquisition mode in positive and negative electrospray ionization as separate analyses, at 1.0 second/scan, 50-1500 Da mass range, and precursor ion collision energy to 6 eV (Function 1, low energy) in trap section with a range from 20-40 eV (Function 2, high voltage) for transfer section. Compounds were pre-identified with Progenesis QI v2.3 for small molecules (Nonlinear Dynamics, Waters Corporation), using Progenesis MetaScope with the Human Metabolome Database (HMDB) v3.5 (Wishart et al., 2012), LIPID MAPS Database (Sud et al., 2006) and ChEBI reference database (Hastings et al., 2012). The search parameters were as follows: precursor tolerance, theoretical fragmentation, and fragment tolerance ≤20 ppm, with isotope similarity ≥90%. Identification parameters were as follows: total score ≥35, fragmentation score ≥90%, and mass error ≤20 ppm (Varela-Rodríguez et al., 2019). Metabolic and statistical analyzes were made with MetaboAnalyst v4.0 (Chong et al., 2018).

For targeted quantification of tricarboxylic acids (TCA), dried extracts were dissolved in a mixture of 5% methanol 95% water with 0.1% formic acid, using 5 µL of solvent per milligram of fresh tissue. A calibration curve of analytical standards was prepared through serial dilutions for analytical standards (Table S2) at 0.1, 0.3, 1, 3, 10, 30 and 100 µM. Each calibration level was analyzed in triplicate and samples in duplicate, injecting 5 µL of each sample. An isocratic method was used for analyte chromatographic separation at 0.3 mL/min flow rate, starting at 1% B during 2.5 min, followed by a wash at 99% B for 1.5 min, and a re-equilibration at 1% B for 2 min. The MS was operated in negative ionization using targeted TOF-MRM mode (Alelyunas et al., 2014). The TOF was operated in sensitivity mode and a resolution of 9,000 FWHM at 0.8 seconds/scan, with the optimized collision energy and target enhanced duty cycle ion used for each analyte area shown in Table S2. Skyline software v4.1 (Henderson et al., 2018; MacLean et al., 2010) was utilized for peak integration and quantification, while statistical analyses were performed with R v3.4.4 (R Core Team, 2018).

## RESULTS

### Identification of *lin-28* and let-7 Factors in Axolotl

Two coding transcripts were reconstructed for potential functional orthologs of *lin-28a* and *lin-28b*, encoded in two genomic loci of the axolotl genome (Nowoshilow et al., 2018). The alignment of translated open reading frames (ORFs) corresponding to amxLin28A and amxLin28B proteins are shown in Fig. S1A. A domain conservation analysis for amxLin28A and amxLin28B denoted the presence of characteristic domains, previously reported for the Lin28 family in other bilaterians (Mayr and Heinemann, 2013). The structural organization of amxLin28A and amxLin28B proteins is conserved, both with a type S1 cold-shock domain (CSD) close to amino-terminal region, a zinc knuckle domain with two retroviral-type zinc fingers (ZnF) near to carboxyl-terminal region, and a predicted nucleolar localization signal (NoLS) between these two domains (Fig. 1A). The distinctive feature among amxLin28A and amxLin28B is the presence of a nuclear localization signal (NLS) in amxLin28B only (Fig. 1A). Through a phylogenetic analysis with *lin-28* coding sequences (CDS) from representative organisms of the subphylum Vertebrata, we noticed that *amx-lin-28a* and *amx-lin-28b* have a close similarity with their respective orthologs (Fig. 1B), especially with amphibian sequences. The alignment of *lin-28* CDS used in the phylogenetic tree for the clade of salamanders is shown in Figure S1B. Moreover, the phylogenetic reconstruction of *lin-28b* sequences seems to recapitulate the phylogeny of vertebrate animals (Maddison et al., 2007; Meyer and Zardoya, 2003). However, the position of *lin-28a* sequences from salamanders and anurans (class Amphibia) suggests an early divergence of these, compared with *lin-28a* sequences from other vertebrates (Fig. 1B).

**Figure 1.**
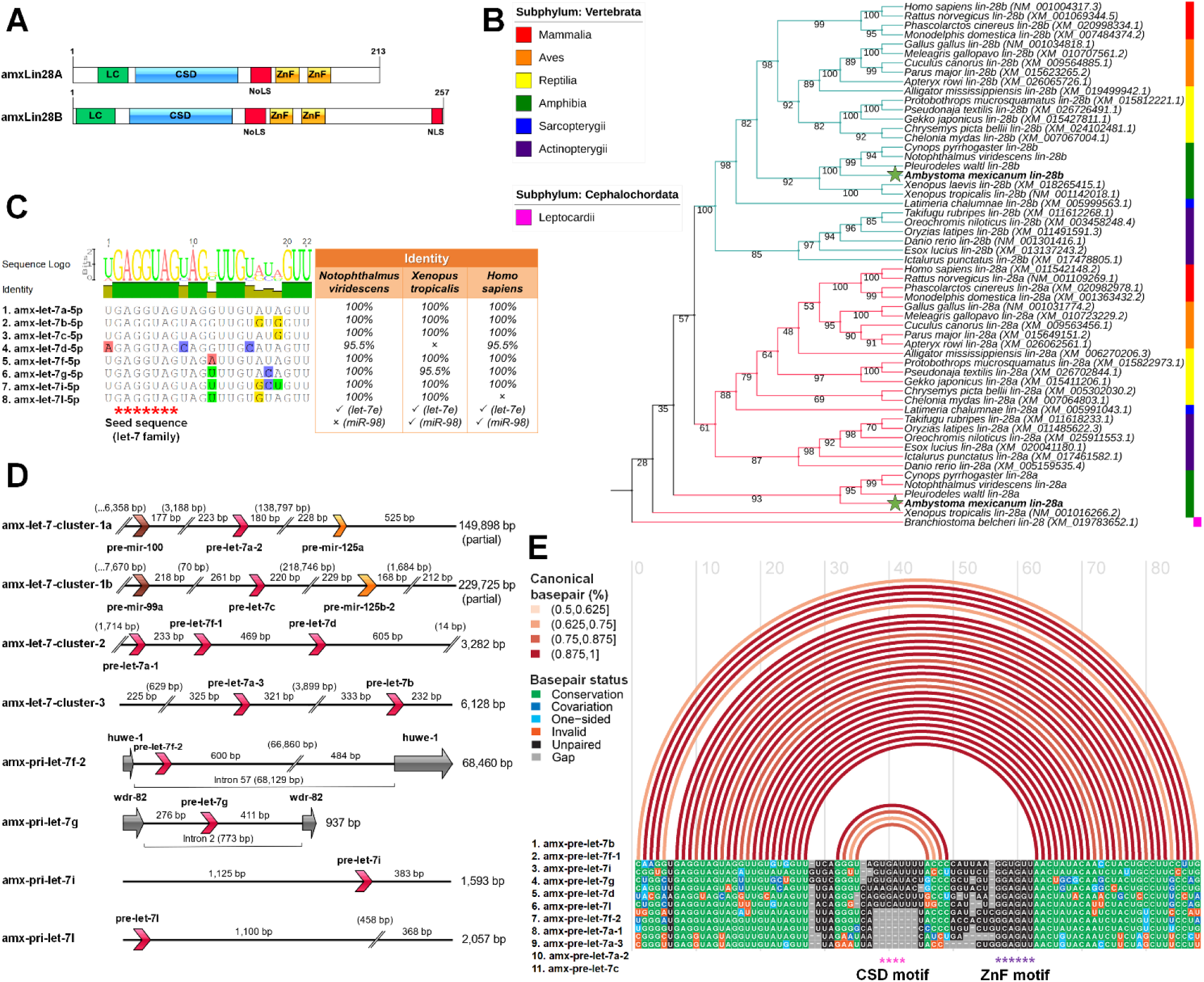
Identification and *in silico* analysis of members for the Lin28 and let-7 families in axolotl. **(A)** Domains organization scheme for canonical sequences translated of the amxLin28A and amxLin28B paralogs, where LC is N-terminal low-complexity region; CSD: S1-like cold-shock domain; NoLS: predicted nucleolar localization signal; ZnF: zinc-knuckle domain comprised by two retroviral-type CCHC zinc-fingers motifs; NLS: C-terminal nuclear localization signal present in amxLin28B only. **(B)** Rooted Maximum Likelihood phylogenetic tree for *lin-28* coding sequences of representative vertebrate animals. Bootstrap support was determined by 10,000 replicates. Branch lengths were omitted for the best visualization of the tree. Branch colors indicate different paralogs of *lin-28* and green stars highlight the *lin-28* sequences of axolotl. **(C)** Mature let-7 microRNAs confirmed in axolotl and their percentage of sequence identity with respect to equivalent members in other tetrapods. Red asterisks mark the seed sequence region corresponding to the let-7 family. **(D)** Genomic organization of primary let-7 transcripts in axolotl. Orientation of the different microRNA precursors and exons are shown with directional arrows in different colors to discriminate between distinct microRNA families (pre-let-7: red arrows; pre-mir-125: orange arrows; pre-mir-100/99: brown arrows), or coding exons (gray arrows). **(E)** Alignment of precursor sequences of the let-7 family in axolotl. Arcs in different shades of red indicate percentages of canonical pairing between base pairs into a secondary folding structure. Asterisks in different colors mark distinct cis-motifs exposed in hairpin-type structures.

In order to find possible homologs of mature let-7 microRNAs in axolotl, we used a sRNA-seq dataset generated from diverse organs (Caballero-Pérez et al., 2018). Thereby, we were able to identify eight different mature microRNAs in axolotl, with an identical seed sequence corresponding to the let-7 family (Fig. 1C). These mature let-7 sequences showed an identity of at least, 95% when compared with their mature let-7 counterparts reported for other tetrapods, evidencing a high conservation from amphibians to mammals. One mature let-7 is only present in amphibians apparently, denominated as let-7l (Fig. 1C). It is noteworthy the absence of a mature let-7e, despite being a widely distributed microRNA in most jawed vertebrates. Also, the lack of mature miR-98 in axolotl and other urodeles revised is notable, since this microRNA is a let-7 family member commonly observed among diverse tetrapods, except in birds. These observations of presence-absence are based on comparisons made with several mature let-7 family sequences reported in the miRBase database v21.0 (Griffiths-Jones et al., 2006).

Through a series of micro-assemblies using a mRNA-seq dataset previously published (Caballero-Pérez et al., 2018), we were able to reassemble eight distinct let-7 transcripts that mapped to eight different loci (Fig. 1D) in the axolotl genome (Nowoshilow et al., 2018). Some primary let-7 transcripts show a tandem clustered arrangement highly conserved across vertebrates (Roush and Slack, 2008). This is the case for the amx-let-7-cluster-1a/b transcripts, whose arrangement involves a mir-10 family precursor (mir-100/99a) near to a pre-let-7, also associated to a second mir-10 family precursor (mir-125) located at a greater distance (Fig. 1D). Other transcripts found as clusters in axolotl, called amx-let-7-cluster-2 and amx-let-7-cluster-3, are also preserved in both the location order and the number of pre-let-7, which only show a distance variation between precursors when compared to their homologs in humans (Fig. 1D). While the let-7-cluster-4 in mammals is constituted by the pre-let-7f-2 and pre-mir-98, the amx-pri-let-7f-2 is contained into the intron 57 of the axolotl *huwe-1* gene without a pre-mir-98 associated (Fig. 1D). This seems to be a common feature between salamanders, given the absence of a mature miR-98 reported for other urodeles, and the fact that we were not able to detect it in any of the sRNA-seq datasets analyzed. The amx-pri-let-7g is also intronically encoded in the axolotl genome (Fig. 1D), with a smaller size in base pairs with respect to its counterpart in humans, but whose localization is maintained within the intron 2 of the axolotl *wdr-82* gene. The remaining primary transcripts cataloged as amx-pri-let-7i and amx-pri-let-7l are monocistronic forms (Fig. 1D). While the amx-pri-let-7i is a conserved paralog in diverse tetrapods, the amx-pri-let-7l seems to be exclusive of amphibians, since the mature let-7l sequence has been only found in some urodeles and anurans revised in this work (Fig. 1C).

Our results show that pri-let-7 transcripts encode for eleven different let-7 precursors (Fig. 1D). A general folding model generated from the secondary structures predicted for these amx-pre-let-7 is presented in Figure 1E. The folding model revealed the typical stem-loop secondary structure for the let-7 family, with the mature sequence localized at stem level and cis-motifs exposed in terminal loops, a structure associated with the recognition and accessibility to Lin28 domains (Nam et al., 2011). It is important to note that the cis-motif sequence necessary for interaction with the cold shock domain of Lin28, is not present in all the amx-pre-let-7 transcripts (Fig. 1E, pink asterisks), while the cis-motif sequence needed for interaction with the ZnF domain of Lin28 is conserved in all amx-pre-let-7 transcripts identified in this study (Fig. 1E, purple asterisks).

### Transcriptional Expression of *lin-28* and let-7 Factors during Forelimb Regeneration

In this work we used late juvenile organisms, which did not manifest any evident sexual dimorphism (Fig. 2A). Each stage across limb regeneration was distinguished by a morphological phenotype, as shown in Figure 2B. The process started from an uninjured limb with a complex composition of tissues and cells in different physiological contexts. After amputation, the first macroscopic response observed was the active generation of a blood clot that stopped the bleeding within a few minutes. Then, an erythematosus thin epithelium was restored during wound healing at 2 days post amputation (dpa). Subsequently, this epithelium became opaque and thick at 6 dpa, when a specialized epithelium called apical epidermal cap (AEC) was forming. The emergence of a recognizable blastema occurred at 14 dpa, revealed by a white cone-shaped protuberance that grew up from the base of amputation plane. Later, the blastema entered to a high cell proliferation phase at 20 dpa, which was denoted by an increased outgrowth of the blastema, maintaining a conical shape but reddened at its base due to an apparent greatest blood irrigation. As limb regeneration continued, morphogenetic events of redifferentiation and patterning were triggered to give rise new organized tissues, establishing a late digital palette until early digits at 42 dpa. The last stage registered at 56 dpa, evoked the maturation of newly formed tissues and subsequent growth of the new structure, which will eventually reach its original size.

**Figure 2.**
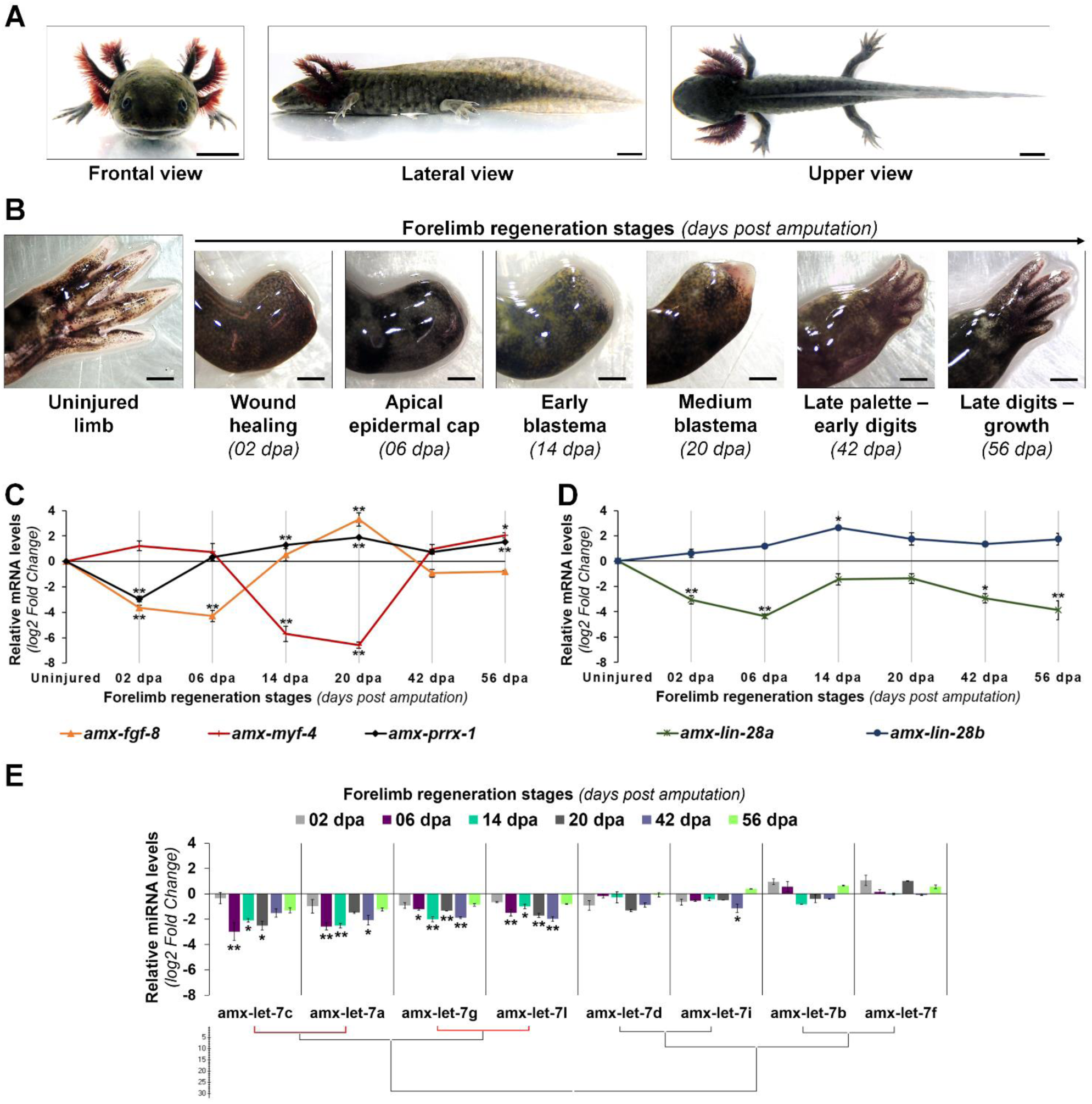
Transcriptional profiles of Lin28 factors and mature let-7 microRNAs during forelimb regeneration. **(A)** Morphological features of a late juvenile axolotl (14-16 cm snout-tail length) from different views. Scale bars: 1 cm. **(B)** Temporal and morphological characterization of the forelimb regeneration stages analyzed in late juvenile axolotls. Scale bars: 1 mm. **(C)** Transcriptional expression profile of some genetic markers during limb epimorphosis. **(D)** Relative abundance of *lin-28* transcripts during epimorphic regeneration. **(E)** Maturation patterns of the let-7 family across regeneration. Hierarchical clustering was made with the Ward algorithm and a Manhattan distance measure, indicating in red a positive Pearson correlation of *P* <0.05. Data in the graphs are represented as mean ± s.e.m. (*n* = 3); *, *P* <0.05; **, *P* <0.01; one-way ANOVA with post-hoc Tukey-Kramer test vs uninjured condition.

In addition, we evaluated the temporal expression profile of some genes whose transcriptional expression has been considered as specific for certain events during epimorphic process (Fig. 2C). The transcript abundance of *amx-prrx-1*, a homeobox factor known as an early marker for blastemal cells (Satoh et al., 2011), displayed a decrease at 2 dpa when compared to the uninjured condition, but it clearly increased in blastema stages with high peaks of expression at 14 and 20 dpa. In the case of the transcript abundance for *amx-fgf-8*, a key inductor for outgrowth of the blastema (Han et al., 2001), a decline was observed at 2 and 6 dpa with a gradually increase at 14 dpa, reaching its highest level of expression at the medium blastema stage (20 dpa). As a marker for cell differentiation, we analyzed the expression pattern of myogenic factor *amx-myf-4*, whose activation has been associated with muscle tissue differentiation (Sandoval-Guzmán et al., 2014). Our results indicate that the transcript levels of *amx-myf-4* drastically decreased at blastema stages (14 and 20 dpa), and gradually increased at 42 and 56 dpa when cellular redifferentiation occurs in epimorphosis (Fig. 2C).

Once the limb regeneration stages for late juvenile axolotls were defined, at morphological and molecular levels, we proceeded to characterize the transcriptional profile of *lin-28* and mature let-7 factors across the process. Expression patterns obtained for both axolotl *lin-28* paralogs were contrasting; while the abundance of *amx-lin-28a* transcripts was downregulated along the process, the *amx-lin-28b* transcripts gradually raised since 2 dpa until blastema stages (Fig. 2D) mainly at 14 dpa. Furthermore, the expression pattern of *amx-lin-28b* is very similar to the previously determined profile for *amx-prrx-1* and coincides with the reported stages of regeneration that displayed an increased cell proliferation (Johnson et al., 2018). Although *amx-lin-28a* transcripts remained low during regeneration, compared with the uninjured limb, some incremental rebounds of transcripts were detected at 14 and 20 dpa. On the other hand, the abundance of most mature let-7 microRNAs showed a gradual decay as epimorphosis progresses, reaching the maximum reduction in blastema stages at 6, 14, and 20 dpa, with a tendency to increase in the subsequent stages at 42 and 56 dpa (Fig. 2E). It should be noted that, even though most mature let-7 microRNAs displayed low abundances during epimorphosis, the mature amx-let-7c, amx-let-7a, amx-let-7g, and amx-let-7l were the most downregulated along the process (Fig. 2E).

Altogether, these results show a generalized downregulation of mature let-7 microRNAs in blastema stages, which present an inverse behavior respect to the transcriptional upregulation of *amx-lin-28b*, suggesting a conservation of the antagonistic functional role for amxLin28B on the maturation of let-7 microRNAs during axolotl limb regeneration.

### amxLin28B shuttles from Cytoplasm to Nucleus during Forelimb Regeneration

According to the human protein atlas (Uhlen et al., 2015), human Lin28 factors are highly expressed in testis and placenta during adult life, leaving aside pathological conditions as cancer. To determine the subcellular localization of axolotl Lin28 proteins, we performed an immunohistochemical analysis with longitudinal cross-sections throughout limb epimorphosis. Particularly, we found that amxLin28B-positive cells (Fig. 3, panels A/C/E/G; Fig. 4, panels I/K/M; red signal) were observed mainly among connective tissue. In uninjured limb (Fig. 3, panel B1), amxLin28B (Fig. 3, panel A1 red signal) was confined to the cytoplasm of dermal fibroblasts (Fig. 3, panel A2 yellow arrow) localized at stratum spongiosum and compactum (Fig. 3, panels B2-3 black arrows), and interstitial fibroblasts (Fig. 3, panel A3 yellow arrow) that resided between muscle fibers (Fig. 3, panels B4-5 black arrows). However, amxLin28B was also detected in some nuclei of epidermal cells, interspersed close to the stratum germinativum and around of Leydig cells (Fig. 3, panel A2 magenta arrow), as well as in some nuclei of muscle fibers (Fig. 3, panel A3 magenta arrow).

**Figure 3.**
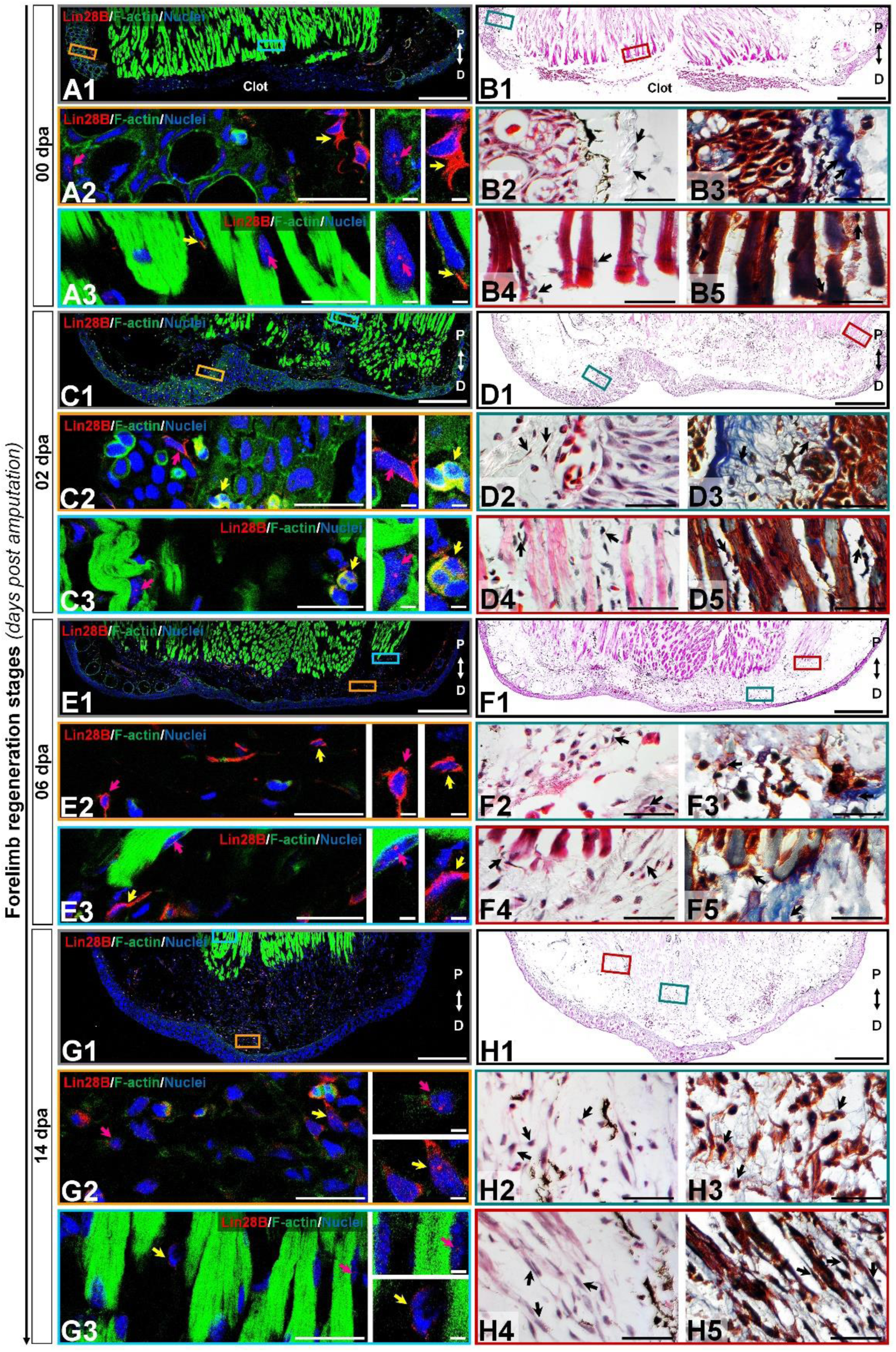
Subcellular immunolocalization of amxLin28B during forelimb regeneration. **(A-H)** Histological sections of representative stages during the limb regeneration oriented in Proximal-Distal axis. **(A-B)** Uninjured limb (00 dpa). **(C-D)** Wound healing stage (02 dpa). **(E-F)** Apical epidermal cap stage (06 dpa). **(G-H)** Early blastema stage (14 dpa). **(A1, C1, E1, G1)** Intracellular localization dynamics of amxLin28B marked in red, by contrast with DAPI (blue) and Phalloidin (green) that delimit nuclei and cell shapes, respectively. Scale bars: 500 µm. **(A2-3, C2-3, E2-3, G2-3)** Magnifications with different color contours to remark some areas of interest for amxLin28B in red, DAPI in blue, and Phalloidin in green; arrows in yellow and magenta have in turn their own zoom in. Scale bars: 50 and 5 µm, respectively. **(B1, D1, F1, H1)** The architectural organization of tissues is shown with Hematoxylin & Eosin staining. Scale bars: 500 µm. **(B2/4, D2/4, F2/4, H2/4)** Magnifications with distinct color contours to indicate some areas of interest in Hematoxylin & Eosin staining. Scale bars: 50 µm. **(B3/5, D3/5, F3/5, H3/5, J3/5, L3/5/7, and N3/5/7)** Histological changes detected on extracellular matrix for similar zones indicated in H&E ampliations, which are displayed with a differential staining to connective tissue. Scale bars: 50 µm. The different arrows indicate diverse tissue components, as described in results.

**Figure 4.**
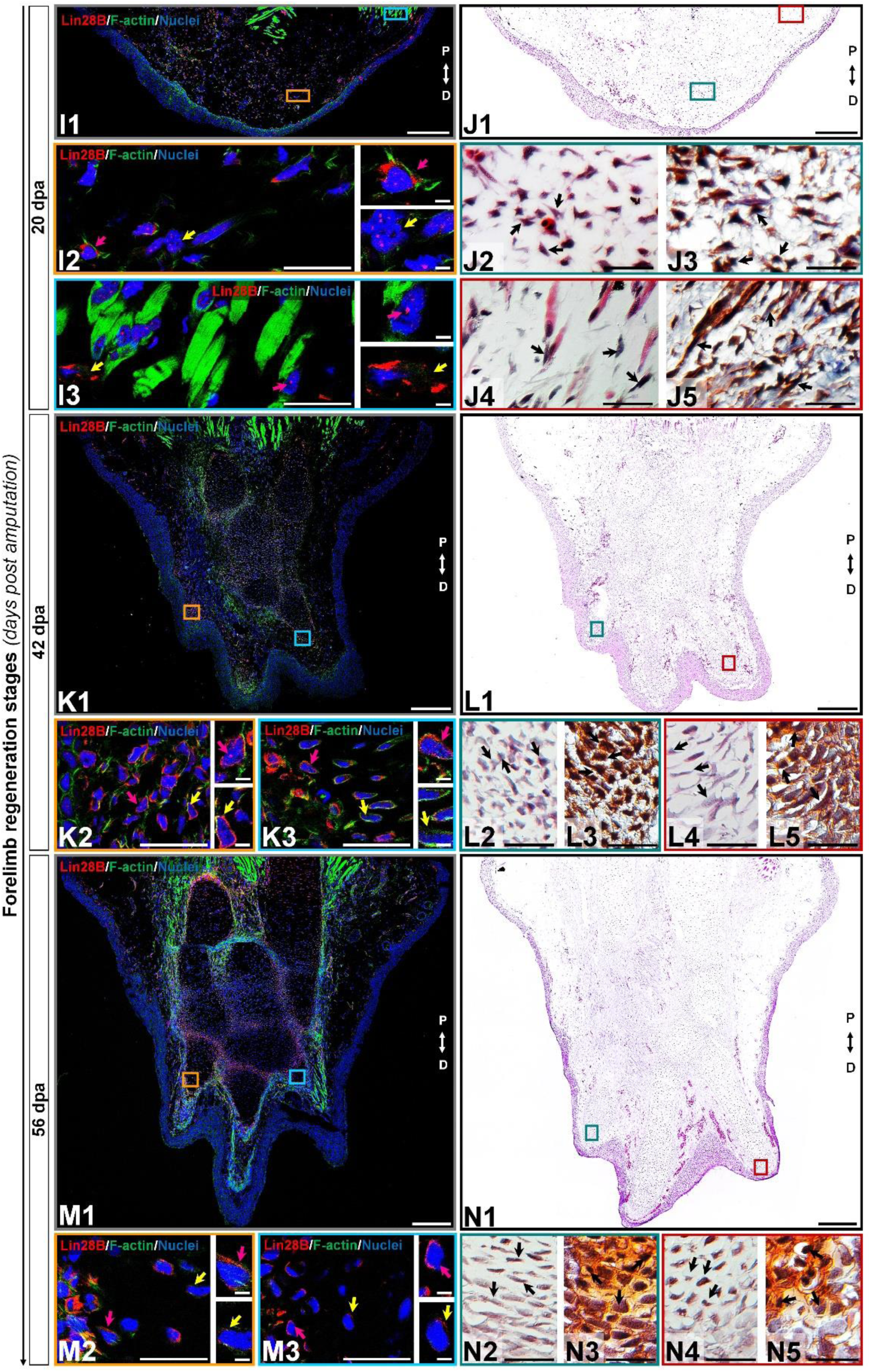
Subcellular immunolocalization of amxLin28B during forelimb regeneration (continuation). **(I-N)** Histological sections of representative stages during the limb regeneration oriented in Proximal-Distal axis. **(I-J)** Medium blastema stage (20 dpa). **(K-L)** Late palette–early digits stage (42 dpa). **(M-N)** Late digits–growth stage (56 dpa). **(I1, K1, M1)** Intracellular localization dynamics of amxLin28B marked in red, by contrast with DAPI (blue) and Phalloidin (green) that delimit nuclei and cell shapes, respectively. Scale bars: 500 µm. **(I2-3, K2-3, M2-3)** Magnifications with different color contours to remark some areas of interest for amxLin28B in red, DAPI in blue, and Phalloidin in green; arrows in yellow and magenta have in turn their own zoom in. Scale bars: 50 and 5 µm, respectively. **(J1, L1, and N1)** The architectural organization of tissues is shown with Hematoxylin & Eosin staining. Scale bars: 500 µm. **(J2/4, L2/4, and N2/4)** Magnifications with distinct color contours to indicate some areas of interest in Hematoxylin & Eosin staining. Scale bars: 50 µm. **(J3/5, L3/5, and N3/5)** Histological changes detected on extracellular matrix for similar zones indicated in H&E ampliations, which are displayed with a differential staining to connective tissue. Scale bars: 50 µm. The different arrows indicate diverse tissue components, as described in results.

**Figure 5.**
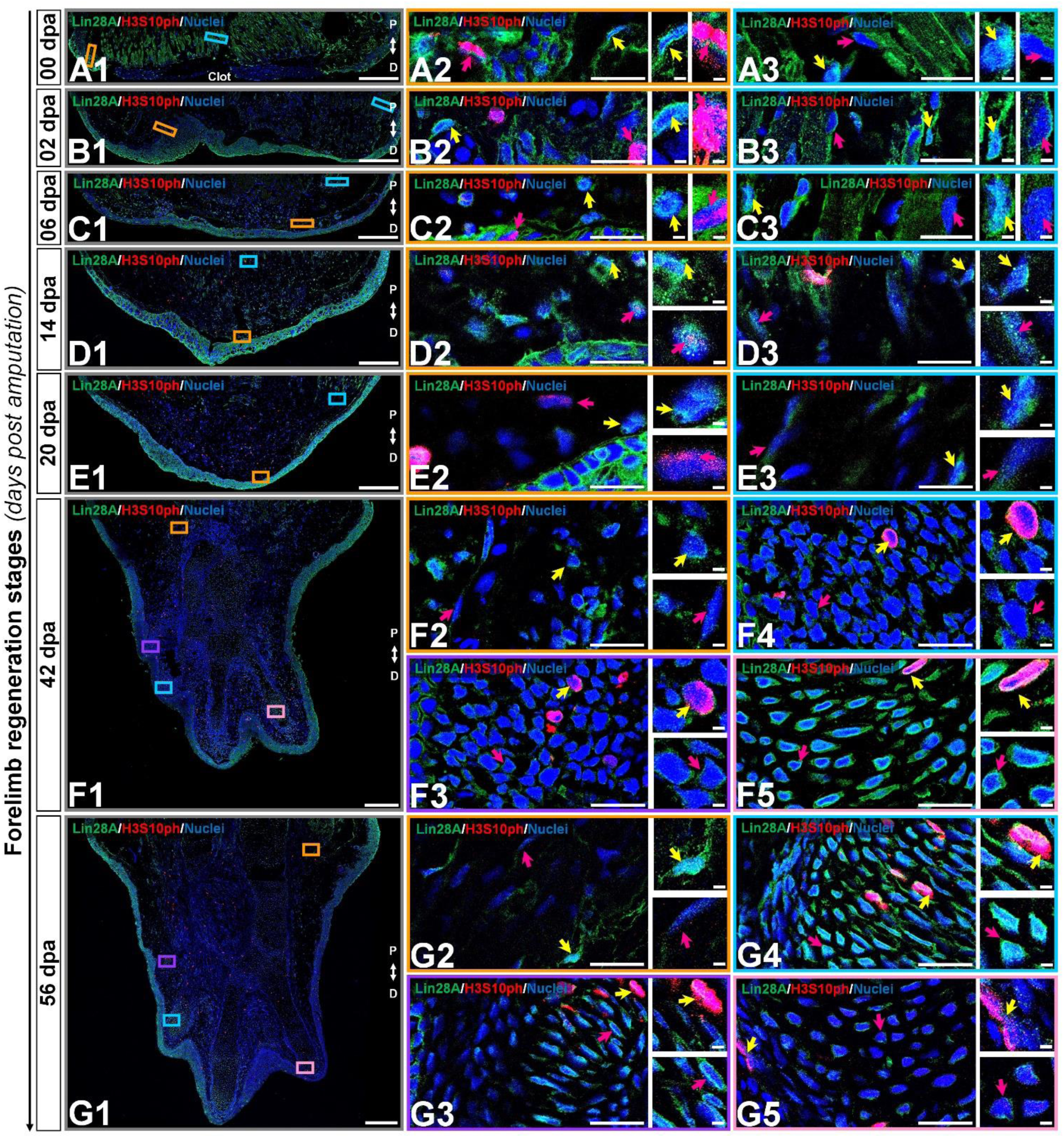
Subcellular immunolocalization of amxLin28A during forelimb regeneration. **(A-G)** Histological sections of key stages during limb epimorphosis arranged in the Proximal-Distal axis. **(A)** Uninjured limb (00 dpa). **(B)** Wound healing stage (02 dpa). **(C)** Apical epidermal cap stage (06 dpa). **(D)** Early blastema stage (14 dpa). **(E)** Medium blastema stage (20 dpa). **(F)** Late palette–early digits stage (42 dpa). **(G)** Late digits–growth stage (56 dpa). **(A1-G1)** Intracellular detection of amxLin28A marked in green, and H3S10ph in red for mitotic cells, by contrast with DAPI (blue) to delimit nuclei. Scale bars: 500 µm. **(A2/3-G2/3)** Ampliations with distinct color contours to distinguish some areas of interest for amxLin28A in green, H3S10ph in red, and DAPI in blue; yellow and magenta arrows have in turn their own magnifications. Scale bars: 50 and 5 µm, respectively. The different arrows mark diverse tissue components, as described in results.

The cytoplasmic localization of amxLin28B in connective tissue was maintained at 2 dpa (Fig. 3, panel D1) and 6 dpa (Fig. 3, panel F1), when re-epithelialization and histolysis occurs, respectively. The amxLin28B-positive cells (Fig. 3, panels C1/E1 red signal) manifested a faithful signal coming from the cytoplasm of dermal fibroblasts (Fig. 3, panels C2/E2 magenta arrows; E2 yellow arrow) associated to collagen fibrils (Fig. 3, panels D2-3/F2-3 black arrows), and interstitial fibroblasts (Fig. 3, panel E3 yellow arrow) embedded among muscle fibers (Fig. 3, panels D4-5/F4-5 black arrows). We noticed that, as previously observed on uninjured limb, some cells continued to display nuclear amxLin28B in epidermis and muscle fibers at 2 dpa (Fig. 3, panel C3 magenta arrow) and 6 dpa (Fig. 3, panel E3 magenta arrow). Nevertheless, a leukocyte infiltration also revealed that amxLin28B is present in the cytoplasm of putative polymorphic nuclear cells (with lobed nuclei), apparently associated with F-actin (Fig. 3, panels C2-3 yellow arrows).

Subcellular localization of amxLin28B changed from the cytoplasm to the nucleus in connective tissue, during the establishment of a blastema at 14 dpa (Fig. 3, panel H1), extending to the proliferative stage of blastema at 20 dpa (Fig. 4, panel J1). Some amxLin28B-positive cells (Fig. 3, panel G1 red signal) showed a transitional state at 14 dpa, such as blastemal cells (Fig. 3, panels H2-3 black arrows) that presented both cytoplasmic and nuclear location of amxLin28B (Fig. 3, panel G2 magenta and yellow arrows). However, amxLin28B at 20 dpa (Fig. 4, panel I1 red signal) was mainly restricted to the nuclei (Fig. 4, panel I2 magenta and yellow arrows) of blastemal cells, which were isolated or forming small aggregates (Fig. 4, panels J2-3 black arrows). In the case of muscle tissue, we detected an active disorganization of the fibers at their apical ends (Figs. 3 and 4, panels H4-5/J4-5 black arrows), while amxLin28B remained nuclear (Figs. 3 and 4, panels G3/I3 magenta arrows). Although several interstitial fibroblasts displayed a cytoplasmic amxLin28B (Figs. 3 and 4, panels G3/I3 yellow arrows), the location of amxLin28B was nuclear in the most distal apical-situated fibroblasts.

When morphogenetic events of redifferentiation and patterning are triggered at 42 dpa (Fig. 4, panel L1), nuclear localization of amxLin28B became mostly cytoplasmic. The amxLin28B signal (Fig. 4, panel K1 red signal) was detected at both nuclear and cytoplasmic levels (Fig. 4, panel K2 magenta arrow) in early digital condensates, constituted in principle by pre-chondrocytes (Fig. 4, panels L2-3 black arrows) derived from putative committed blastemal cells, and even some amxLin28B-positive cells only presented cytoplasmic amxLin28B (Fig. 4, panel K2 yellow arrow). In the case of late digital condensates, outermost cells showed predominantly cytoplasmic amxLin28B (Fig. 4, panel K3 magenta arrow), while innermost cells maintained this behavior but with tendency to decrease the amxLin28B signal (Fig. 4, panel K3 yellow arrow). In addition, we also noted that more mature chondrocytes started to form rudimental cartilaginous scaffolds, revealed as orange areas stained by safranin (Fig. 4, panels L4-5 black arrows).

Finally, a miniature limb has been formed at 56 dpa (Fig. 4, panel N1), showing a localization of amxLin28B similar to that observed in the uninjured limb. Peripheral amxLin28B-positive cells (Fig. 4, panel M1 red signal) of perichondrium exhibited cytoplasmic signal (Fig. 4, panels M2-3 magenta arrows), which gradually decreased as internalization and maturation of chondrocytes occurs (Fig. 4, panels M2-3 yellow arrows). Additionally, most preaxial digit presented an advanced chondrogenesis with trabeculae and a dense cartilage matrix, stained in light orange with safranin (Fig. 4, panels N4-5 black arrows), while postaxial digits had fewer trabeculae (Fig. 4, panels N2-3 black arrows), according to a pre-to-postaxial gradient of differentiation.

Altogether, these results evidence a highly dynamic subcellular localization for amxLin28B throughout regeneration. Alternation of amxLin28B, from cytoplasm to nucleus, happens during recruitment and proliferation of blastemal cells mainly at 14 and 20 dpa. In contrast, the nuclear to cytoplasmic mobilization of amxLin28B occurs during early chondrogenic redifferentiation process at 42 dpa.

### amxLin28A remains in the Cell Cytoplasm during Forelimb Regeneration

In the case of amxLin28A, we found that amxLin28A-positive cells (Fig. 5, panels A-G green signal) were prominently viewed among epidermal tissue. In the uninjured limb, and from 2 to 6 dpa, amxLin28A (Fig. 5, panels A1/B1/C1 green signal) was detected in the cytoplasm of epidermal cells with mitotic H3S10ph-positive cells (Fig. 5, panels A2/B2/C2 magenta arrows), putative pericytes (Fig. 5, panel B2 yellow arrow), diverse fibroblasts (Fig. 5, panels A2-3/B3/C2-3 yellow arrows), muscle fibers (Fig. 5, panels A3/B3/C3 magenta arrows), and glandular tissue. However, a high amxLin28A signal was reached in the cytoplasm of epidermal cells at 14 and 20 dpa (Fig. 5, panels D1/E1 green signal), with mitotic H3S10ph-positive cells (Fig. 5, panels D2/E2 magenta arrows) noticed between blastemal cells. Particularly, multiple blastemal cells showed presumptive cytoplasmic structures of rods and rings with amxLin28A at 14 dpa (Fig. 5, panel D2 yellow arrow), which were not recognized at 20 dpa (Fig 5, panel E2 yellow arrow). Likewise, amxLin28A was confined to the cytoplasm of muscle fibers (Fig. 5, panels D3/C3 magenta arrows) and fibroblasts (Fig. 5, panels D3/E3 yellow arrows) at 14 and 20 dpa.

On the other hand, digital condensates established at 42 and 56 dpa showed amxLin28A (Fig. 5, panels F1/G1 green signal) in the cytoplasm of putative pre-chondrocytes (Fig. 5, panels F3-4 magenta arrows). Although chondrocytes with active synthesis of cartilaginous matrix also exhibited cytoplasmic amxLin28A from 42 dpa, it was mainly perceived around of nuclear periphery (Fig. 5, panels F5/G3/G4 magenta arrows), in contrast to the chondrocytes of most developed preaxial digital element (Fig 5, panel G5 magenta arrow) with a dense cartilage scaffold at 56 dpa. Some mitotic H3S10ph-positive cells were concentrated around of digital condensates (Fig. 5, panels F3-4 yellow arrows), as well as in the prospective perichondrium and its surroundings (Fig. 5, panels F5/G3-5 yellow arrows), where a niche of chondroprogenitor cells appears to be maintained. For muscle fibers and fibroblasts, amxLin28A was retained in the cytoplasm (Fig. 5, panels F2/G2 magenta and yellow arrows) at 42 and 56 dpa.

### Effects of the Functional Lin28 Inhibition on the Forelimb Regeneration

The compound Lin28-1632 is a dose-dependent chemical inhibitor for interactions among Lin28 proteins and its targets. The inhibitory action of Lin28-1632 on Lin28 proteins has been successfully tested *in vitro* using murine embryonic stem cells (Roos et al., 2016) and *in vivo* with an allotransplanted mouse mammary carcinoma (Chen et al., 2019).

By topical administration of Lin28-1632, we questioned its inhibitory effect on axolotl Lin28 factors at specific regeneration stages, following the treatment scheme shown in Figure 6A (see Methods for details). At 6 dpa, only a relatively thin epidermis was noted covering the amputation site in limbs treated with inhibitor, compared to the controls (vehicle and without treatment) (Fig. 6B). At 14 dpa, a smaller cone-shaped blastema was perceived in limbs treated with the inhibitor, compared to the controls (Fig. 6B). Most notable phenotypic alteration was evidenced at 20 dpa, where a smaller blastema was viewed in limbs treated with inhibitor, compared to the controls (Fig. 6B). At 42 dpa, limbs treated with inhibitor were slightly smaller with defined digital elements, and a less interdigital space between postaxial digits, contrasting with the controls (Fig. 6B). Morphometric analysis of the photographic record revealed a significant reduction of regenerated areas after treatments with inhibitor at 14, 20, and 42 dpa, showing a difference of ∼40% for blastema stages compared to the control conditions (Fig. 6C). These findings suggest that the topical application of Lin28-1632 could cause a decay of cell proliferation and/or an increase of differentiation, due to the functional inhibition of Lin28.

**Figure 6.**
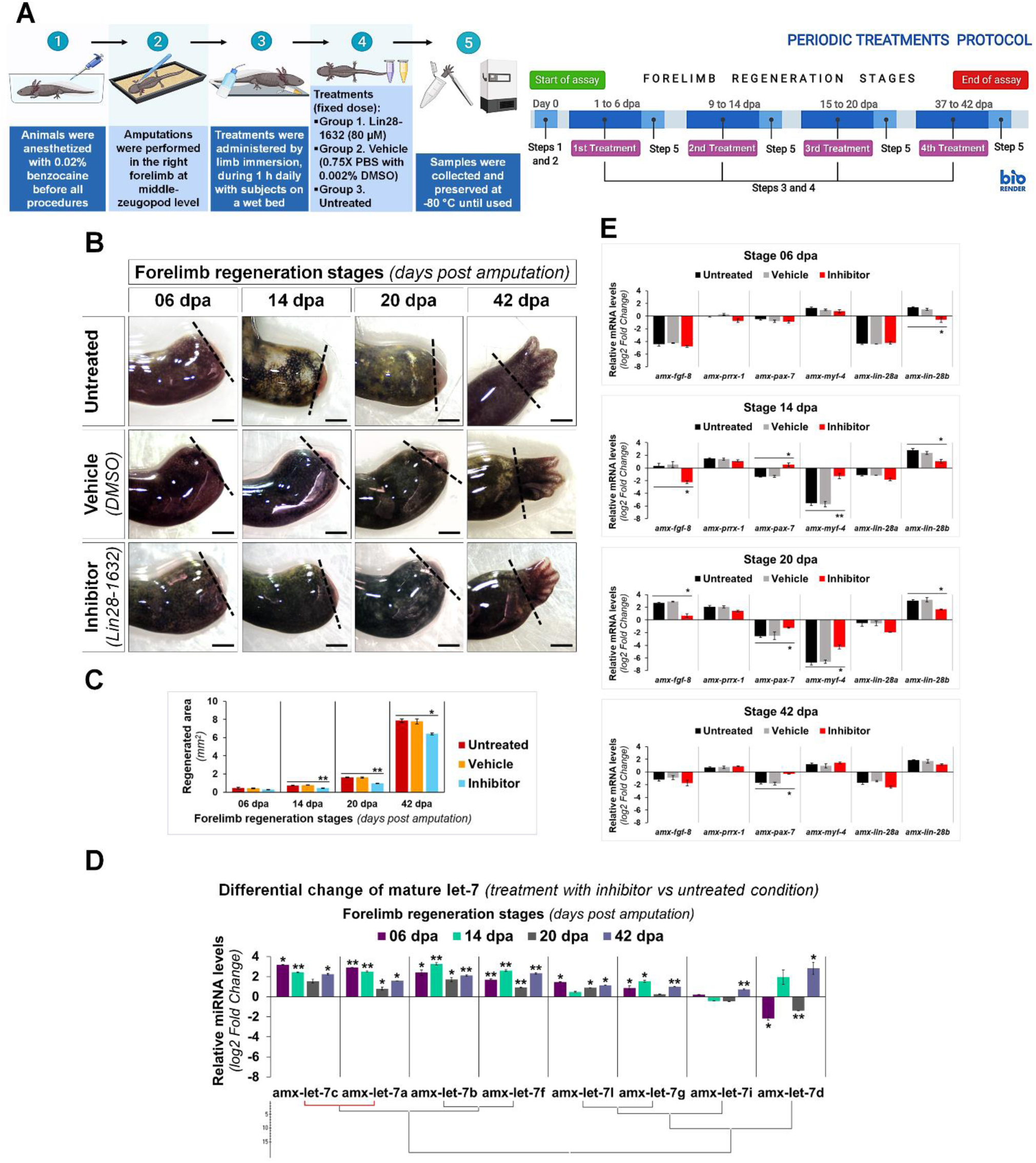
Characterization of induced effects after Lin28 inhibition in key regeneration stages. **(A)** Schematic diagram of the periodic administration protocol followed; on the left side, it shows the general strategy implemented, and on the right side, the temporary management schedule to each regeneration stage of interest. Created with BioRender. **(B)** Morphological changes observed after administration of different treatments in most representative stages of regeneration. Dotted lines indicate the amputation site. Scale bars: 1 mm. **(C)** Morphometric analysis of regenerated areas to conditions assayed. **(D)** Differential change in maturation patterns of the let-7 family in representative regeneration stages after treatments with inhibitor (vs respective stages untreated). Hierarchical clustering of Ward with Manhattan distance measurement, marking in red a positive Pearson correlation of *P* <0.05. **(E)** Relative transcript abundances of some genetic markers during key stages of limb epimorphosis. Data in the graphs are represented as mean ± s.e.m. (*n* = 3); *, *P* <0.05; **, *P* <0.01; one-way ANOVA with post-hoc Dunnett test vs untreated control condition.

In order to explore the implications of the functional inhibition of Lin28 factors after Lin28-1632 treatments (Fig. 6A, see Methods for details), we measured the relative levels of let-7 microRNAs and transcripts for some marker genes. Many of the mature let-7 microRNAs significantly increased their abundance after pharmacological inhibition of Lin28 at different stages (Fig. 6D). Among these microRNAs, the most upregulated were amx-let-7c, amx-let-7a, amx-let-7b, and amx-let-7f. At 6 dpa, only a significant reduction of *amx-lin-28b* transcripts was detected in samples treated with inhibitor, compared to the controls (Fig. 6E). At 14 and 20 dpa, the transcript levels for *amx-lin-28b* and *amx-fgf-8* significantly decreased in samples treated with inhibitor, compared to the controls (Fig. 6E). On the other hand, the abundance of *amx-myf-4* transcripts was drastically reduced in the controls at 14 and 20 dpa, while the samples treated with inhibitor showed a significantly less pronounced decrease (Fig. 6E). A similar behavior was observed for *amx-pax-7* transcripts, whose abundance significantly increased after treatments with inhibitor at 14, 20, and 42 dpa, compared to the controls (Fig. 6E). Although changes in transcript levels for *amx-lin-28a* and *amx-prrx-1* were not statistically significant, the abundance of *amx-lin-28a* transcripts tended to decrease in the blastema stages of inhibitor-treated samples (Fig. 6E).

Since previous morphological analysis evidenced changes in the limbs treated with Lin28-1632, we also examined the histological alterations after Lin28-1632 treatments (Fig. 6A, see Methods for details). At 6 dpa, the absence of a clear apical space between muscle fibers and epidermis was noted in samples treated with inhibitor (Fig. 7, panels C1/D1), while a blastema began to develop in the control (Fig. 7, panels A1/B1). Likewise, the inhibitor-treated limb sections displayed fibroblasts (Fig. 7, panels D2-3 black arrows) and putative pericytes with cytoplasmic amxLin28B (Fig. 7, panel C2 magenta arrow) similarly to the control (Fig. 7, panel A2 magenta and yellow arrow), although various dermal fibroblasts were not strongly associated to collagen fibrils in the control condition (Fig. 7, panels B2-3 black arrows), consistent with the recruitment of blastemal cells. Moreover, a greater leukocyte infiltration was observed in samples treated with inhibitor (Fig. 7, panels C2-3 yellow arrows) compared to the control (Fig. 7, panel A3 yellow arrow), revealed by the presence of putative polymorphic nuclear cells (with lobed nuclei and yellow signal). In the case of muscle fibers, nuclear localization of amxLin28B was maintained in both sample conditions, treated with inhibitor (Fig. 7, panel C3 magenta arrow) and vehicle (Fig. 7, panel A3 magenta arrow). However, a general integrity of muscle fibers was detected for treatments with inhibitor, especially at their apical ends (Fig. 7, panels D4-5 black arrows), contrasting with muscle fibers of the control with loose apical ends (Fig. 7, panels B4-5 black arrows), relative scarce endomysium, and some cells released from them.

**Figure 7.**
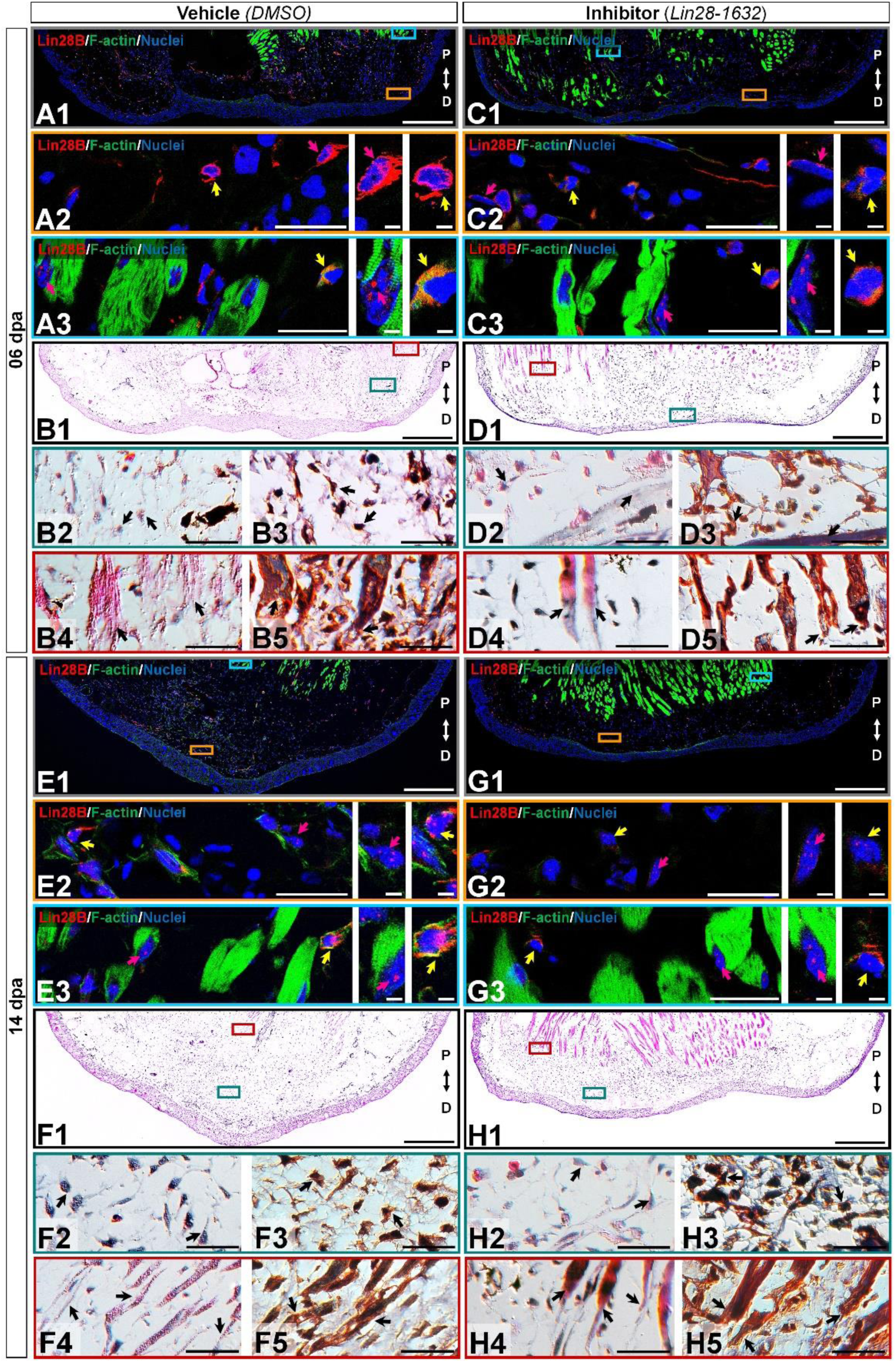
Histological changes after functional inhibition of Lin28 in key regeneration stages. **(C, D, G, H)** Histological sections of limbs treated with Lin28-1632 inhibitor, oriented in Proximal-Distal axis. **(A, B, E, F)** Histological sections of limbs treated with vehicle, oriented in Proximal-Distal axis. **(A-D)** Apical epidermal cap stage (06 dpa). **(E-H)** Early blastema stage (14 dpa). **(A1, C1, E1, G1)** Subcellular localization of amxLin28B marked in red, by contrast with DAPI (blue) and Phalloidin (green). Scale bars: 500 µm. **(A2-3, C2-3, E2-3, G2-3)** Magnifications with different color contours to remark some areas of interest for amxLin28B in red, DAPI in blue, and Phalloidin in green; arrows in yellow and magenta have in turn their own zoom in. Scale bars: 50 and 5 µm, respectively. **(B1, D1, F1, H1)** General tissue morphology shown with Hematoxylin & Eosin staining. Scale bars: 500 µm. **(B2/4, D2/4, F2/4, H2/4)** Magnifications with distinct color contours to indicate some areas of interest in Hematoxylin & Eosin staining. Scale bars: 50 µm. **(B3/5, D3/5, F3/5, H3/5)** Differential staining to view changes on stroma for similar zones indicated in H&E ampliations. Scale bars: 50 µm. The distinct arrows point diverse tissue components, as described in results.

At 14 dpa, the samples treated with inhibitor (Fig. 7, panels G1/H1) presented a smaller blastema with a thin apical epidermal cap, compared to the control (Fig. 7, panels E1/F1). In the inhibitor-treated limb sections, blastema tissue showed few cells with amxLin28B signal localized both in the cytoplasm and the nucleus (Fig. 7, panel G2 magenta and yellow arrows). Furthermore, a low cell density was observed, along with abundant fibrin and collagen fibrils (Fig. 7, panels H2-3 black arrows) in the inhibitor-treated samples. These features contrast with those viewed in the control condition, whose blastemal cell density was higher (Fig. 7, panels F2-3 black arrows) with several cells exhibiting amxLin28B signal, both in cytoplasmic and nuclear compartments (Fig. 7, panel E2 yellow arrow), or in the nucleus only (Fig. 7, panel E2 magenta arrow). We also noticed that, as previously described for treatments in the regeneration stage at 6 dpa, muscle fibers continued to display nuclear amxLin28B in samples treated with inhibitor (Fig. 7, panel G3 magenta arrow), similarly to the control (Fig. 7, panel E3 magenta arrow). Likewise, interstitial fibroblasts exhibited cytoplasmic amxLin28B, both in vehicle and inhibitor treatments (Fig. 7, panels E3/G3 yellow arrows). However, muscle fibers of the vehicle-treated control presented a greater disorganization mainly at their apical ends (Fig. 7, panels F4-5 black arrows), while muscle fibers of inhibitor-treated samples maintained a certain compaction and integrity (Fig. 7, panels H4-5 black arrows) with few cells released from them.

At 20 dpa, tissue sections of limbs treated with inhibitor showed a thin epidermis and a smaller blastema (Fig. 8, panels K1-L1) than the control (Fig. 8, panels I1-J1). Few blastemal cells presented cytoplasmic amxLin28B (Fig. 8, panel K2 magenta and yellow arrows) in the inhibitor-treated samples, noticing very few cells with faint signs of nuclear amxLin28B, as well as a low cell density (Fig. 8, panels L2-3 black arrows). Conversely, the vehicle-treated control exhibited a dense population of blastemal cells (Fig. 8, panels J2-3 black arrows) with both nuclear and cytoplasmic amxLin28B signal (Fig. 8, panel I2 yellow arrow), or only nuclear (Fig. 8, panel I2 magenta arrow). In the case of muscle fibers, both inhibitor and control treated samples manifested nuclear amxLin28B (Fig. 8, panels I3/K3 magenta arrows) with amxLin28B-positive interstitial fibroblasts having cytoplasmic signal (Fig. 8, panels I3/K3 yellow arrows). Nonetheless, muscle fibers of inhibitor-treated limbs revealed more compressed distal ends (Fig. 8, panels L4-5 black arrows), while in the control condition was detected a greater disorganization of muscle fibers with loose distal ends (Fig. 8, panels J4-5 black arrows).

**Figure 8.**
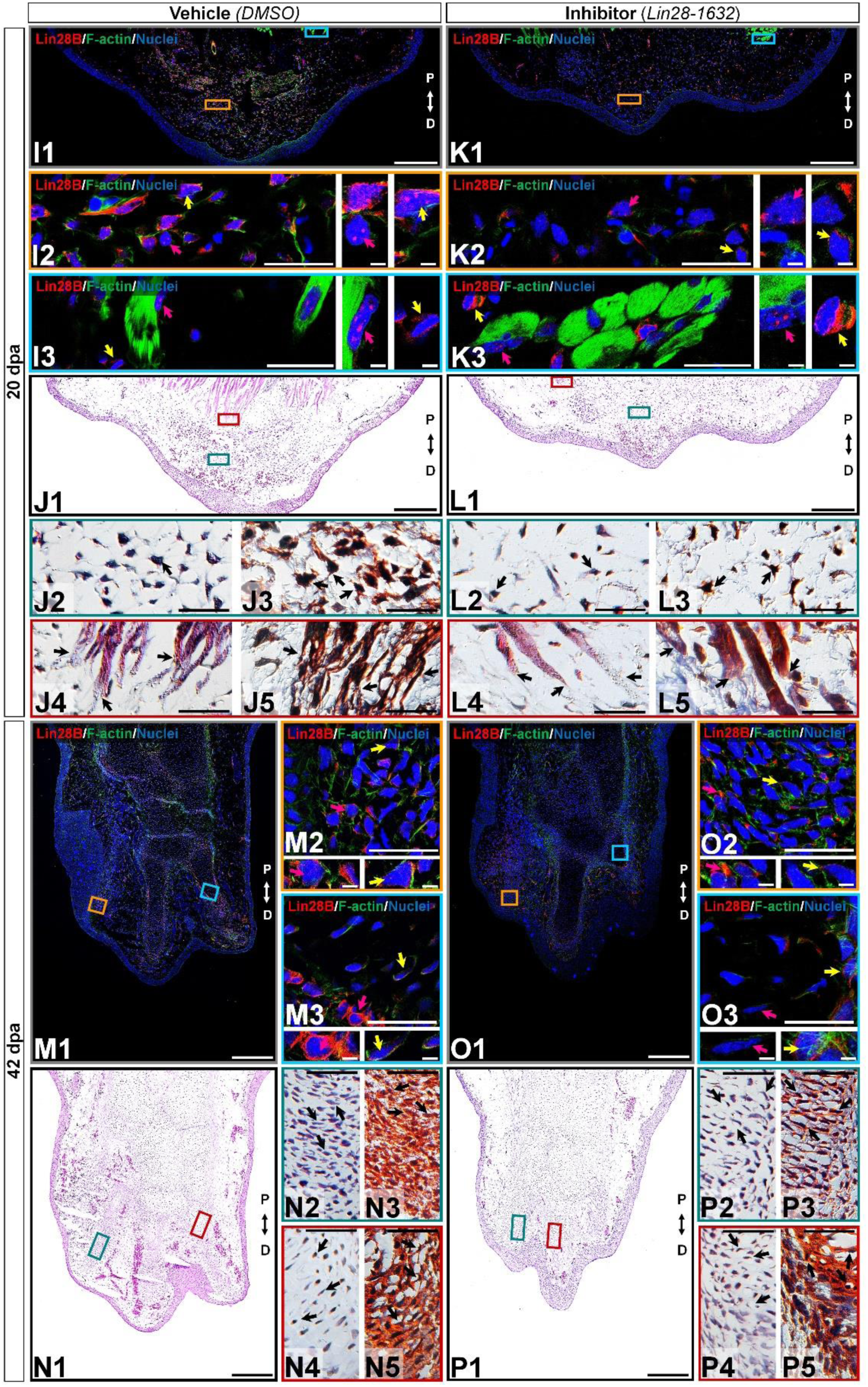
Histological changes after functional inhibition of Lin28 in key regeneration stages (continuation). **(K, L, O, P)** Histological sections of limbs treated with Lin28-1632 inhibitor, oriented in Proximal-Distal axis. **(I, K, M, N)** Histological sections of limbs treated with vehicle, oriented in Proximal-Distal axis. **(I-L)** Medium blastema stage (20 dpa). **(M-P)** Late palette–early digits stage (42 dpa). **(I1, K1, M1, O1)** Subcellular localization of amxLin28B marked in red, by contrast with DAPI (blue) and Phalloidin (green). Scale bars: 500 µm. **(I2-3, K2-3, M2-3, O2-3)** Magnifications with different color contours to remark some areas of interest for amxLin28B in red, DAPI in blue, and Phalloidin in green; arrows in yellow and magenta have in turn their own zoom in. Scale bars: 50 and 5 µm, respectively. **(J1, L1, N1, P1)** General tissue morphology shown with Hematoxylin & Eosin staining. Scale bars: 500 µm. **(J2/4, L2/4, N2/4, P2/4)** Magnifications with distinct color contours to indicate some areas of interest in Hematoxylin & Eosin staining. Scale bars: 50 µm. **(J3/5, L3/5, N3/5, P3/5)** Differential staining to view changes on stroma for similar zones indicated in H&E amplifications. Scale bars: 50 µm. The distinct arrows point diverse tissue components, as described in results.

At 42 dpa, a relatively small limb was observed in samples treated with inhibitor (Fig. 8, panels O1-P1), compared to the control (Fig. 8, panels M1-N1). Postaxial digital condensates of inhibitor-treated limb sections exhibited cells with cytoplasmic amxLin28B (Fig. 8, panel O2 magenta arrow), mainly perceived as a faint signal for cells concentrated inside the chondrogenic aggregates (Fig. 8, panel O2 yellow arrow). On the other hand, the vehicle-treated control showed cells with preferentially cytoplasmic amxLin28B (Fig. 8, panel M2 magenta arrow), and very few cells with nuclear traces of amxLin28B (Fig. 8, panel M2 yellow arrow). It is worth noting that Lin28-1632 treatments induced a shorter interdigital space among postaxial digital condensates, with premature synthesis of collagen (blue stained) and cartilage (orange stained) (Fig. 8, panel P3 black arrows). Such putative premature chondrocytes seem to display an atypical organization that resembles the trabeculae observed in advanced differentiation stages (Fig. 8, panel P2 black arrow), contrasting with the control (Fig. 8, panels N2-3 black arrows). The most preaxial digital element of the inhibitor-treated samples revealed some chondrocytes with cytoplasmic amxLin28B in the perichondrium (Fig. 8, panel O3 yellow arrow), and around the internal digital periphery (Fig. 8, panel O3 magenta arrow). In contrast, we noticed clear cytoplasmic amxLin28B signal for perichondrium cells (Fig. 8, panel M3 magenta arrow), and some cells near to internal digital periphery (Fig. 8, panel M3 yellow arrow), in the vehicle-treated control. Moreover, cartilage deposition (orange stained) was pronounced in the most preaxial digital scaffold of samples treated with inhibitor (Fig. 8, panels P6-7 dark arrows), having several visible trabeculae and a perichondrium with collagen (blue stained). While some internal trabeculae can be seen in the most preaxial digital element of the control condition (Fig. 8, panels N4-5 black arrows), inner peripheral areas in close contact with the perichondrium had less cartilage (light orange stained).

Although the pharmacological inhibition of Lin28 altered the proper regeneration process, leading to an aberrant developmental phenotype, the regenerative event still occurs. Therefore, we questioned if a prolonged treatment with the Lin28-1632 compound could drastically affect the regeneration process. In this way, a continuous treatment scheme was implemented, whose details are described in Figure S2A (see Methods for details). The sustained application of inhibitor caused the formation of a premature smaller limb with syndactyly and four rudimentary digits at 56 dpa (Fig. S2B), compared to the controls. Even though continuous application of Lin28 inhibitor did not fully inhibit the regenerative process, we observed at histological level a less vascularization of the limb with shorter digital cartilaginous elements, and without evident joints (Fig. S2C), contrasting with the control conditions. These phenotypic alterations after sustained Lin28 inhibition revealed more drastic changes during epimorphosis than those previously described (Figs. 6-8), suggesting that pharmacological inhibition of amxLin28 proteins by Lin28-1632 might be affecting the process in a time and dose-dependent manner, as previously tested (Chen et al., 2019; Roos et al., 2016).

Altogether, these results indicate that functional inhibition of amxLin28 proteins through the Lin28-1632 administration alters the proper regeneration process, attributable in part to upregulation of some let-7 microRNAs and the consequent repression of their targets, such as the Lin28 family. However, it is not ruled out that other direct target transcripts for Lin28 factors could be also affected.

### Metabolic Reprogramming during Forelimb Regeneration

Previous studies have involved the Lin28/let-7 circuit in modulating primary cell metabolism (Shyh-Chang et al., 2013). To explore the role of the circuit on potential metabolic reprogramming events during epimorphosis, we scrutinized global metabolic changes in the process employing an UPLC-HDMS. Using a Sparse Partial Least Squares-Discriminant Analysis (sPLS-DA), the samples were classified according to each stage of regeneration, clustering together the biological replicates (Fig. 9A). This method allowed a clear discrimination of the samples belonging to stages 20, 42, and 56 dpa, which are well separated from other groups. In the case of samples corresponding to the stages 6 and 14 dpa, as well as samples for uninjured limb and 2 dpa, showed an overlapping of their respective confidence regions attributable to a lower diversity in the abundance of shared metabolites and, therefore, a greater similarity between these stages. An agglomerative clustering confirms the similarities observed between samples at metabolic level, forming three main clusters (Fig 9B). The majority of biological replicates for 20, 42 and 56 dpa stages have more similarities between them and constituted one cluster, while a second cluster is formed by samples of 2, 6 and 14 dpa, associated to a third cluster conformed for samples of uninjured condition with some replicates of 2 and 56 dpa.

**Figure 9.**
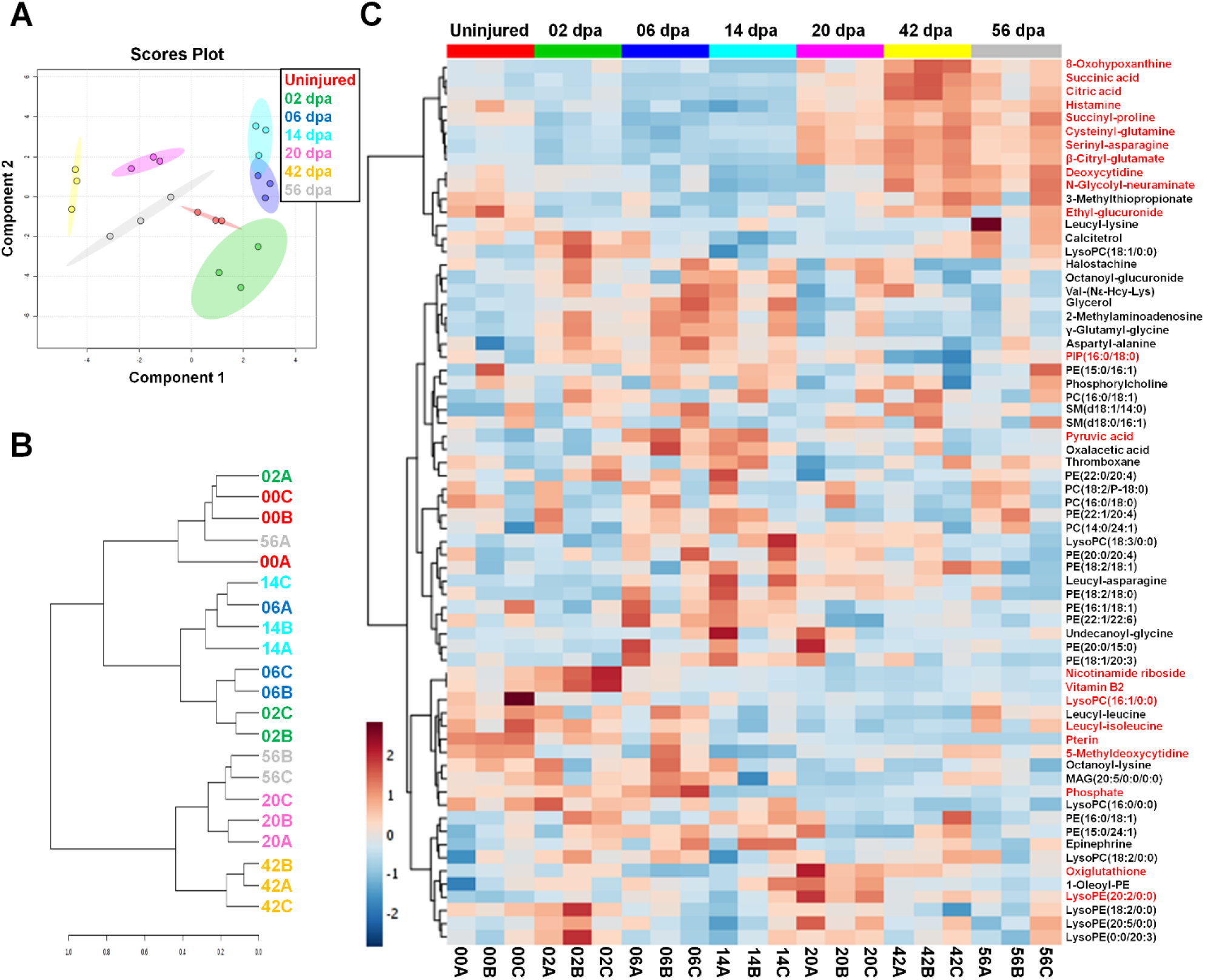
Metabolic profiling during the regeneration process. **(A)** Clustering of metabolite samples evaluated by Sparse Partial Least Squares-Discriminant Analysis (sPLS-DA) with 5-fold Cross-Validated threshold. The stages are highlighted in different colors, displaying regions with a 95% confidence for samples grouped. **(B)** Hierarchical clustering dendrogram for metabolite samples analyzed through a Spearman distance measure and Ward clustering linkage. The stages are shown with different colors. **(C)** Heatmap of normalized abundances for pre-identified metabolites during regeneration. The metabolite abundances highlighted in red have a *P* <0.05; one-way ANOVA with post-hoc Tukey-Kramer test vs uninjured condition. Hierarchical clustering of metabolites was made using a Euclidean distance measure and the Ward clustering algorithm.

The global metabolic profiling made across axolotl regeneration was plotted as a heatmap (Fig. 9C). The non-targeted metabolite analysis revealed 67 compounds that comprises the metabolic diversity detected, including highly hydrophilic metabolites such as organic acids; metabolites of intermediate polarity, for instance, some dipeptides that were considered as presumptive breakdown products of protein catabolism according to the HMDB (Wishart et al., 2012); and hydrophobic metabolites, such as miscellaneous structural lipids, among other metabolites. The complete list of pre-identified metabolites and their corresponding relative abundances are shown in Table S3. Particularly, an increased anabolism appears to occur during establishment of an early blastema at 6 and 14 dpa, due to the high abundance of different classes of phospholipids such as phosphatidylcholine (PC) and phosphatidylethanolamine (PE), with a low abundance of several dipeptides (Fig. 9C). This anabolic state in turn may be related with a favoring of glycolytic pathway at expense of the Krebs cycle (TCA cycle) since a greater abundance of pyruvic acid was found with a less abundance of citric and succinic acids. In this way, an increased catabolism seems to take place during regeneration stages with a high cell proliferation (20 dpa), early re-differentiation (42 dpa), and tissue maturation/latent growth (56 dpa). This catabolic activity was evidenced by a progressive increase in TCA cycle intermediates, as citric and succinic acids, accompanied by a significant depletion of pyruvic acid and various phospholipids, while a large proportion of dipeptides rich in proline, glutamine, asparagine, and serine also increased (Fig. 9C). In addition, it should be noted a substantial abundance of lyso-type lipids at 2 and 20 dpa, such as lysophosphatidylethanolamine (lysoPE) and lysophosphatidylcholine (lysoPC), when a significant cell migration befalls during active inflammation and blastemal cell proliferation, respectively.

In order to understand the implications of metabolic changes occurred at key stages of regeneration, we performed a metabolite set enrichment analysis coupled to a pathway analysis in contrast to the uninjured limb (Fig. 10A). At 6 dpa, it was appreciated an overrepresentation of glutamate, arachidonic acid, and glutathione pathways (Fig. 10A), probably related to the declining inflammation that persists according to previous histological analyzes. Some metabolites with the potential to exacerbate a proinflammatory state were downregulated, such as citric acid and histamine, which could be associated with a fine modulation reported of the inflammatory process (Godwin et al., 2013). These findings are consistent with a moderate abundance observed of thromboxane, also described as an inflammatory modulator (Capra et al., 2014). In addition, the low abundance of oxiglutathione may be indicative of less availability in reduced glutathione, due to the high abundance detected of gamma-glutamyl-glycine, since a sustained gamma-glutamyl transferase activity has been reported for this regeneration stage (Stewart et al., 2013).

**Figure 10.**
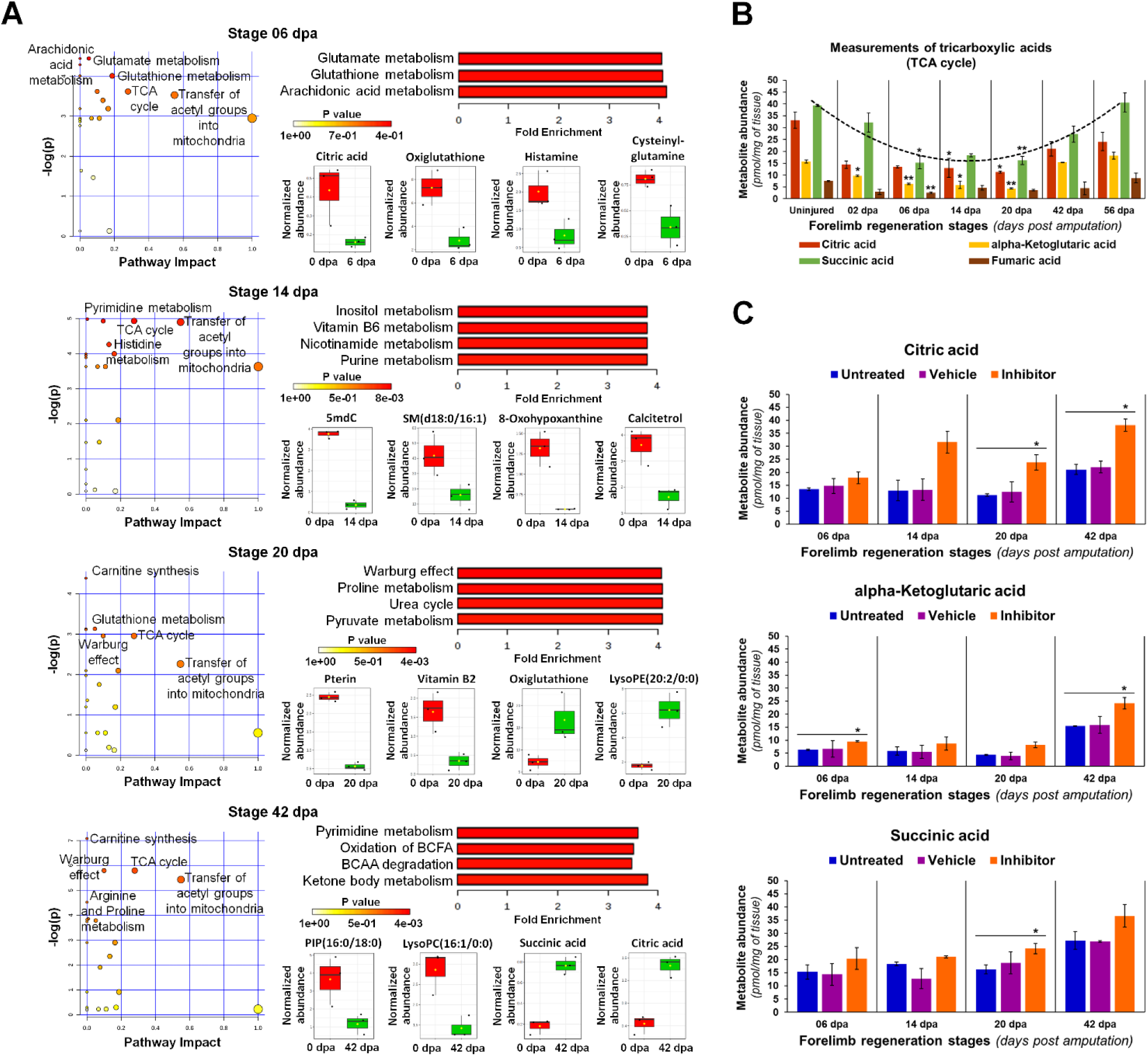
Metabolic analyses in key regeneration stages. **(A)** Pathway analysis by global network topology evaluation with relative-betweenness centrality measure to estimate the importance of nodes. Different dot sizes represent the matched pathway status. Quantitative Metabolite Set Enrichment Analysis (QMSEA) is shown next with metabolite sets more enriched only, using human pathway-associated metabolite sets SMPDB as reference through a global test. The box and whisker plots display some representative metabolites with a significant differential abundance (2-fold change threshold, *P* <0.05 with unpaired t-Student test vs uninjured condition). BCFA: branched-chain fatty acids; BCAA: branched-chain amino acids. **(B)** Abundance validation of some tricarboxylic acids (TCA). Illustrative trend line for succinic acid is shown in a dotted line (*R*^2^ >0.9). Data in the bar plot are represented as mean ± s.e.m. (*n* = 3); *, *P* <0.05; one-way ANOVA with post-hoc Tukey-Kramer test vs uninjured condition. **(C)** Quantification of some tricarboxylic acids after diverse treatments. Inhibitor was the Lin28-1632, and the vehicle 0.75X PBS with 0.002% DMSO. Data in the bar plots are represented as mean ± s.e.m. (*n* = 3); *, *P* <0.05; one-way ANOVA with post-hoc Dunnett test vs untreated control condition.

At 14 dpa, most enriched pathways are associated to the metabolism of purine and pyrimidine nucleotides, such as vitamin B6, nicotinamide, and histidine (Fig. 10A). In this context, the low significant abundance viewed of hypoxanthine seems to indicate a downregulation of purine nucleotide catabolism (Papandreou et al., 2019). Several classes of PE and PC phospholipids increased their abundance, while diverse dipeptides of leucine, lysine, asparagine, serine, proline, and glutamine presented low levels. However, the production of sphingomyelins was reduced despite the abundance of PC, which is a known precursor substrate to ceramides of sphingolipids. In the case of calcitetrol, an inactive form of calcitriol (Christakos et al., 2015), its abundance decreased at blastema stages (6 to 20 dpa) mainly at 14 dpa, being consistent with an increase of osteoclast activity during bone resorption previously observed at histological level. A similar trend was also noticed for the abundance detected of 5-methyldeoxycytidine.

At 20 dpa, overrepresentation of proline and pyruvate metabolism highlights a presumptive Warburg effect (Fig. 10A) (DeBerardinis and Chandel, 2016; Phang et al., 2015). An increased catabolism of proteins was noted, according to the abundance of varied dipeptides, which explain the overrepresentation of pathways such as urea cycle and ammonia recycling. The significant increase of oxiglutathione suggests a major production of reactive oxygen species (ROS), which may be derived from certain activity of the TCA cycle, since citric and succinic acids increased with depletion of vitamin B2. According to these results, blastemal cells seem to favor aerobic glycolysis (Warburg effect), together with some oxidation of fatty acids and carnitine synthesis, considering that pyruvate and several phospholipids presented a moderate to low abundance (Melone et al., 2018). In addition, significant low levels of pterin were also detected, while other types of phospholipids were abundant, such as lysoPE.

At 42 dpa, most enriched pathways are related to cell energetic metabolism, as the catabolism of branched-chain fatty acids (BCFA) and branched-chain amino acids (BCAA), suggesting an increased mitochondrial activity (Fig. 10A) (Clemót et al., 2020; Neinast et al., 2019). Active lipolysis of phospholipids and triacylglycerols, reflected in the low abundance of several BCFA species included lysoPC and lysoPE, explain the promotion of ketone body metabolism and the Krebs cycle (TCA) (Grabacka et al., 2016). In this sense, the synthesis of carnitine is required to regulate and transfer acyl and acetyl groups into mitochondria, in the form of acyl-carnitine and acetyl-carnitine, respectively (Melone et al., 2018), resulting in the increase of some TCA cycle metabolites, as observed for succinic and citric acids. On the other hand, some dipeptides rich in valine, leucine, and isoleucine, presented a low abundance, while those composed of arginine and proline were abundant, revealing a possible differential metabolism for BCAA. Moreover, the abundance of pyrimidines also increased, such as deoxycytidine and 5-methyldeoxycytidine (Fig. 9C).

Previous described results indicate a central role for the Krebs cycle and transfer of acetyl groups into mitochondria. Therefore, we performed a targeted quantification to some tricarboxylic acids along regeneration (Fig. 10B). Most significant changes were observed in blastema stages (6 to 20 dpa), where organic acids such as citric, succinic, and alpha-ketoglutaric, presented a low abundance when compared to the uninjured limb. The general trend noticed was a high abundance of the TCA cycle metabolites measured at regeneration stages with a high degree of cell differentiation; while the stages with a highest proportion of proliferative and less differentiated blastemal cells displayed low levels of these organic acids (Fig. 10B, black dotted line). To determine if the Lin28/let-7 circuit acts upstream of this primary metabolic behavior during regeneration, we administered the Lin28-1632 compound to induce a functional inhibition of Lin28 factors, following the periodic treatment scheme described in Fig. 6A (See Methods for details). Our results revealed significant changes in the abundance of some organic acids for samples treated with inhibitor, increasing the levels of citric acid at 20 and 42 dpa, alpha-ketoglutaric acid at 6 and 42 dpa, and succinic acid at 20 dpa, when compared to the controls (Fig. 10C).

Overall, our metabolic profiles show a significant reduction of some TCA cycle metabolites at blastema stages (6 to 20 dpa), which increase again at differentiation stages (42 to 56 dpa). In addition, complementary metabolic analyses coupled to the functional Lin28 inhibition, suggest that direct or indirect targets of Lin28 factors and mature let-7 microRNAs, may influence primary metabolic pathways. Therefore, the Lin28/let-7 circuit has the potential to modulate the cell metabolism during axolotl regeneration, which in turn, can impact on cellular energy and the availability of precursor substrates for biological processes which are pivotal for the proper formation of a new limb.

## DISCUSSION

The Lin28/let-7 circuit has been conserved since the appearance of bilateral symmetry in metazoans, although a significant gene number expansion for the Lin28 and let-7 families has occurred among vertebrates throughout evolution (Moss and Tang, 2003). The presence of two Lin28 paralogs is common in vertebrates, while the number of let-7 precursors is variable and tends to progressively increase along the animal phylogeny (Hertel et al., 2012). In this study, we report the identification of Lin28A and Lin28B orthologs in axolotl, each of them showing a structural domain conservation with respect to their homologs in humans. Also, we found eleven let-7 precursors encoded in the axolotl genome. Particularly, the let-7 precursors of axolotl seem to group into two sub-classes: one group of pre-let-7 with a clear cis-motif for the cold-shock domain of Lin28 (CSD+), and a second group without such cis-motif (CSD-), similarly to the grouping reported for the twelve human let-7 precursors (Ustianenko et al., 2018). In addition, the cis-motive for the zinc-knuckle domain (ZnF) of Lin28 is highly conserved at sequence level in most pre-let-7 transcripts of axolotl. Previous studies have shown that an interaction with the ZnF is required and sufficient to induce the oligouridylation and degradation of pre-let-7 (Wang et al., 2017), indicating that axolotl let-7 precursors are potentially regulated by amxLin28 proteins, although with a variable affinity for each let-7 member/group.

The function of the Lin28/let-7 circuit has not been directly studied in the context of axolotl regeneration, but previous works have shown an increase of Lin28 protein levels as epimorphosis progresses (Rao et al. 2009), while several let-7 microRNAs undergo a significant downregulation at blastema stages (Holman et al., 2012; King and Yin, 2016). These observations are consistent with our findings, where the establishment and subsequent gradual expansion of a blastemal cell population is accompanied by a progressive increase in *amx-lin-28b* transcripts, along with a generalized continuous decay of the mature amx-let-7 microRNAs. In addition, amxLin28B is expressed in connective tissue and blastemal cells during epimorphosis, and although it has a nuclear localization signal, is not immediately translocated to the nucleus in fibroblasts-like cells. Indeed, subcellular mobilization at specific times and tissues, constitutes an integral part for some intrinsic and extrinsic factors. This may be the case for amxLin28B, whose variable subcellular localization could allow a fine-tuning of cellular reprogramming of fibroblast-like cells to blastemal cells during epimorphosis. In accordance with this proposition, Merlin protein has previously been shown to act as a cell density-dependent growth suppressor in mammalian Schwann cells, through interaction with Lin28B. When cell density is high, Merlin is dephosphorylated to sequester Lin28B, by anchoring it to the actin cytoskeleton, resulting in the maturation of certain microRNAs as the let-7 family, and the subsequent inhibition of cell proliferation. Conversely, when cells lose cell-to-cell contacts, Merlin is phosphorylated and releases Lin28B, which then localizes in the nucleus and exerts its down-regulating role on mature let-7 levels (Hikasa et al., 2016).

Moreover, when amxLin28 proteins are functionally repressed by Lin28-1632 occurs an increase of most amx-let-7 microRNAs, coupled to a decrease in *amx-lin-28b* transcript levels. Among the most affected mature let-7, throughout the regenerative process and in response to treatments with the inhibitor, were identified amx-let-7c and amx-let-7a. These mature let-7 derive from precursors that lack a cis-motif for interaction with the CSD (amx-pre-let-7c/a-1/a-2/a-3), indicating that the interaction between Lin28 and pre-let-7 during regeneration is mainly based on the action exerted through the ZnF domain, as reported in other models (Guo et al., 2006; Wang et al., 2017). Since the *amx-lin-28a* transcript levels do not change dramatically after Lin28-1632 treatments, it is possible that the phenotypic abnormalities observed with mis-regulation of amx-let-7 microRNAs are caused by a functional downregulation of amxLin28B. This points to a double-negative feedback loop among amxLin28B and the amx-let-7 microRNAs in axolotl, and suggests that the complementary sequence present in *amx-lin-28b* transcripts might be a functional target for the let-7 family (data not show). In mammals, such double-negative feedback regulation of the Lin28/let-7 circuit has been well characterized, where Lin28 proteins repress the biogenesis of let-7 precursors, and in turn, mature let-7 microRNAs bind to 3’-UTR of Lin28 transcripts to affect its translation (Rybak et al., 2008). On the other hand, the decrease of *amx-fgf-8* transcripts observed in the Lin28 inhibition experiments, is consistent with findings reported during the gastrulation process of *Xenopus*, where FGF and NodaI/Activin pathways were compromised after Lin28 knockdown (Faas et al., 2013). Furthermore, this downregulation may also explain the small size observed in regenerated areas and interdigital spaces after the periodic treatments with Lin28-1632, since a sustained expression of Fgf8 is associated with loss of interdigital cell death and increased cell proliferation, generating syndactyly during mouse limb development (Bosch et al., 2018). Likewise, the TGF-β pathway can be negatively regulated by let-7 microRNAs through repression of type I receptors in *Xenopus* (Colas et al., 2012), which may be related with the premature chondrocyte differentiation reported in this study after Lin28 inhibition, revealed by a shortening of digital elements and enhanced cartilage deposition. In agreement with these findings, during chicken limb development has been reported that the TGFβ pathway sustains the skeletal connective tissue outgrowth, coordinating the formation of chondrogenic aggregates, fibrogenesis, and the correct development of tendons (Lorda-Diez et al., 2009).

In the particular case of skeletal muscle tissue, we note that amxLin28B is confined to the cell nucleus, while amxLin28A is localized at the cytoplasm, even before the regenerative process begins in axolotl. These observations are consistent with the Lin28 detection in differentiated mouse muscle tissue, both cardiac and skeletal, where Lin28 binds to polysomes and stress granules in translation initiation complexes to improve the translational efficiency of various factors, such as Igf2 (Polesskaya et al., 2007). However, certain levels of mature let-7 are required to promote the later stages of muscle differentiation through down-regulation of diverse let-7 targets, such as Hmga2, Dicer1, and Igf2bp1 (a repressor of Igf2 translation), along with the increase in Myogenin expression (Kallen et al., 2013). When limbs are treated with the Lin28-1632 inhibitor, the transcript levels of *amx-pax-7* and *amx-myf-4* are upregulated in different regeneration stages, together with a substantial increase of mature let-7 levels. These factors are important determinants of myogenic lineage, while the Myogenin transcripts and mature let-7 microRNAs are quite downregulated at blastema stages during epimorphosis, suggesting that blastemal cell population in the inhibitor-treated limbs either maintain partially muscle identity, or enter prematurely to cell differentiation. In this sense, it has been shown that the proliferation capacity of muscle cells, and the regeneration failure in aged muscles, are modulated by let-7 action (Drummond et al., 2011). The increase of *amx-pax-7* transcripts after Lin28-1632 treatments could represent a response to a differentiation-inducing stimulus, as occurs in human oculopharyngeal muscular dystrophy, where satellite cells enter to an early proliferative arrest with overexpression of Pax7, MyoG, and Hmgb1, also accompanied by high levels of mature let-7 (Cappelletti et al., 2019). In turn, the increase of *amx-myf-4* transcripts after Lin28 inhibition also supports the notion of an induced muscle differentiation, since the disorganization of muscle fibers was affected, a necessary event for mobilization of satellite cells and interstitial fibroblast cells. These findings are congruent with the relevance of a long-term partial dedifferentiation for remnant-muscle ends during axolotl epimorphosis, which also facilitates the subsequent reconnection between new and old muscle fibers (Wu et al., 2015).

Several studies have linked the Lin28/let-7 circuit with changes in cell metabolism, modulating pathways such as IGF-PI3K-mTOR (Wu et al., 2015). Indeed, some targets of the Lin28/let-7 circuit can act directly on key checkpoints of primary metabolism (glycolysis and TCA cycle), such as Pdk1 (Ma et al., 2014), Hk2 (Docherty et al., 2016), and Pfkp (Shyh-Chang et al., 2013). In this manner, perturbations on the Lin28/let-7 circuit could affect the balance between anabolic and catabolic reactions during axolotl limb regeneration. An overview of the global metabolic variations experienced across epimorphosis, evokes an enhanced biosynthetic activity at early blastema stages (6 and 14 dpa), revealed by the accumulation of metabolic precursors such as pyruvate, oxaloacetate, and diverse phospholipids, necessary to sustain cell proliferation and growth. This anabolic state has been shown to be consistent with a downregulation of let-7 microRNAs, and the upregulation reported for diverse proteins related to cell transcription and translation, mRNA processing, and cell protection by chaperones, which reflect an increased protein synthesis during early blastema stages (Rao et al. 2009). Furthermore, it has also been reported a differential expression of transcripts related to cell growth control, osteoclast activity, and deoxyribonucleotide production before DNA synthesis (Voss et al., 2015).

Interestingly, the global metabolic profiling highlights a presumptive Warburg effect (aerobic glycolysis) at 20 dpa, revealed by a moderate to low abundance of pyruvate and phospholipids, moderate levels of dipeptides with proline and glutamine amino acids, as well as increased levels of succinic and citric acids. These findings closely resemble the metabolic behavior observed in malignant neoplastic cells that, in addition to the high glycolysis rates, suffer a metabolic rewiring to sustain the production of ATP by anaplerotic pathways, such as glutaminolysis and β-oxidation of lipids (DeBerardinis and Chandel, 2016; Melone et al., 2018; Phang et al., 2015). In this context, such metabolic environment might improve cell proliferation during axolotl regeneration, keeping a balance between consumption and production of substrates intended for cell energy and survival. Moreover, we speculate that lyso-type phospholipids could also have a potential role, altering intracellular calcium fluxes through G-protein-coupled receptors (GPR), and thus promote cell migration and proliferation as described for mammal tumorigenic cells (Drzazga et al., 2017; Lee et al., 2017). On the other hand, late stages of the limb regeneration present an enhanced catabolism, evidenced by low levels of several phospholipids and a high abundance of various dipeptides. These observations are consistent with a previous study reporting the upregulation of transcripts related to phospholipids, lipoproteins, and fatty acids, revealing an active metabolism of steroids and cholesterol (Voss et al., 2015). In fact, this catabolic state might be necessary to supply enough acetyl-CoA for ATP production, and thus satisfy the cellular energy demand during differentiation events, since our analysis detected an increase of certain TCA cycle intermediary metabolites, such as succinic and citric acids.

The quantification of some TCA cycle tricarboxylic acids during axolotl epimorphosis reveals a significant depletion for such metabolites mainly at blastema stages, when let-7 microRNAs are low and the *amx-lin-28b* transcripts are abundant. These observations are consistent with a downregulation reported for enzymes of the TCA cycle and mitochondrial electron transport at blastema stages (Rao et al. 2009; Sibai et al., 2019). Likewise, inverted abundance patterns of some TCA cycle metabolites quantified at blastema stages after Lin28-1632 treatments, suggests that the Lin28/let-7 circuit acts upstream of cell reprogramming events related to the TCA cycle. This agrees with the findings made in mouse PSCs, where knockout cells to *Lin-28b-/-*displayed an enriched expression signature for “Mitochondrial inner membrane”, “NADH dehydrogenase activity”, and “Oxidative phosphorylation”, coupled to an increase of TCA cycle intermediaries such as alpha-ketoglutarate, succinate, isocitrate, and oxaloacetate (Zhang et al., 2016).

In summary, our findings indicate the relevance of a fine regulation mediated by the Lin28/let-7 circuit, mainly for metabolic rewiring events that occur during the establishment and expansion of the blastema. In this context, proliferative blastemal cells seem to induce a reminiscent metabolism of the Warburg effect, where a differential maturation of the let-7 microRNAs in turn may represent a conserved molecular mechanism to fine-tune a global dose of the let-7 family and, in this way, generate a more favorable cellular environment for regeneration. Although a detailed tissue-specific characterization of the complex metabolic reprogramming that takes place across axolotl epimorphosis would provide more information in this regard, our results are consistent with previous observations. In fact, cellular behavior can be modified in concrete ways through modifications in metabolism, as happens during immune cell activation or cellular transformation, where some metabolites can act as inductors of a given cellular state in addition to its canonical role (Ganeshan and Chawla, 2014; Ryan et al., 2019). Thus, new opportunities are provided to design promising therapeutic modalities based on metabolic strategies, which could potentiate the regeneration in other organisms with limited capacity. Future approaches in this sense should consider targets of the amxLin28 factors and amx-let-7 microRNAs as candidates to scrutinize specific molecular mechanisms, taking into consideration that the balance between cell proliferation and differentiation also involves a metabolic reprogramming.

## ETHICS STATEMENT

The animal care and procedures carried out were previously approved by the Institutional Animal Care and Use Committee of CINVESTAV (IACUC protocol number 0209-16), following standard practices in accordance with the “guidelines for use of live amphibians and reptiles in field and laboratory research” by the American Society of Ichthyologists and Herpetologists in USA. Likewise, statutes of the Official Mexican Norm NOM-062-ZOO-1999 for the “technical specifications for the production, care and use of laboratory animals” were fulfilled, which are based on the “Guide for the Care and Use of Laboratory Animals” “The Guide” by the National Research Council of USA. Federal Registration Number #B00.02.01.01.01.0576/2019 awarded by the Secretariat of Agriculture and Rural Development (SADER) in México.

## AUTHOR CONTRIBUTIONS

Conceived the project: ACR and HVR. Designed the experiments: HVR and ACR. Performed the experiments: HVR, DGAQ, LVR, DGZ, AEC, and JJOO. Analyzed the data: HVR, LVR, DGZ, JCP, JJOO, and ACR. Wrote the paper: HVR and ACR, with inputs from all co-authors.

## FUNDING

This work was supported by the Swedish International Research Links Grant 2014-9040-114152-32, Consejo Nacional de Ciencia y Tecnología grants: Fronteras de la Ciencia CONACyT FOINS-301 and Ciencia Básica CB-2015-252126. HVR was supported by CONACyT fellowship 247353. DGZ was supported by CONACyT postdoctoral fellowship.

## DATA AVAILABILITY

New Reported sequences and metabolites generated during the current study will be available once the manuscript is accepted for publication in a regular journal.

Supplementary figures, tables, (cited along the text) and other data will be available once this manuscript is accepted.

## CONFLICT OF INTEREST

The authors have no conflict of interest to declare.

## ACKNOWLEDGEMENTS

The authors thank M.C. Gilberto Marquez-Chavoya for technical assistance with the histological procedures. Dr. María Cruz-Santos for initial concepts about the Lin28/let-7 circuit. Biol. Arturo Vergara-Iglesias from CIBAC-UAM Xochimilco and Biol. Ollin O. Ramírez-Sánchez from PIMVS Ambystomania, for the support with the animal sampling and management. We also thank Dr. Marco A. Leyva-González for critically reviewing this manuscript.

## Notes

### Competing Interest Statement

The authors have declared no competing interest.

